# A Drug-Elicitable Alternative-Splicing Module (DreAM) for Tunable AAV Expression and Controlled Myocardial Regeneration

**DOI:** 10.1101/2024.07.01.601517

**Authors:** Zhan Chen, Luzi Yang, Yueyang Zhang, Yue Li, Gonglie Chen, Ze Wang, Jiting Li, Yuhan Yang, Dongyu Zhao, William T Pu, Ke Yang, Erdan Dong, Yuxuan Guo

## Abstract

Adeno-associated virus (AAV) is a major vector for gene therapy. A technique to fine-tune the time and level of AAV expression is lacking, which greatly restricts indication choice, therapeutic efficacy and safety. To solve this problem, here we developed a drug-elicitable alternative-splicing module (DreAM) responsive to risdiplam, an FDA-approved alternative splicing modulator. Risdiplam activates DreAM-regulated AAV expression in a dose-dependent manner with an over 2000-fold inducible change depending on the inducer dosage and the organ of interests. With a temporally resolution of a couple of days, DreAM could repeatedly activate AAV expression, determined by the frequency and duration of risdiplam administration. As an example of DreAM-realized gene therapy, this technique was used to control AAV-based cardiac delivery of YAP^5SA^, a potent cardiomyocyte regeneration factor. DreAM transiently activated AAV-YAP^5SA^, establishing the dedifferentiation-proliferation-redifferentiation cycle in cardiomyocytes. Consequently, DreAM fulfilled the regenerative capacity of AAV-YAP^5SA^ in myocardial infarction while circumventing toxicity associated with prolonged AAV-YAP^5SA^ expression. Together, these data demonstrated a tremendous potential of DreAM to enhance the efficacy, safety and applicable scope of gene therapy.

## Introduction

An optimal gene therapy strategy requires the therapeutic transgene to be expressed at the correct location, time and level. These key parameters are heavily determined by the gene delivery vectors. For example, adeno-associated virus (AAV) transduces a broad spectrum of organs such as heart, skeletal muscle and liver^1^. Proper selection of tissue-specific promoters and AAV capsids could achieve targeted AAV gene expression in these organs^2^. AAV gene expression could last for years, which is suitable for gene supplementation therapy^3^. However, AAV lacks the ability to flexibly adjust the duration and level of transgene expression, which profoundly restricts their applicable indications and raises additional concerns about efficacy and safety, especially in scenarios when transient or fluctuated gene expression is more desirable.

Technology for gene expression regulation has been quickly evolving. Classic techniques like the tet-on/off system^4^ are based on transcriptional regulation, composing a small-molecule inducer, an engineered transcription factor and a special promoter carrying inducer-responsive cis-regulatory elements. The large size and the complicated components of these techniques preclude effective delivery by AAV. Lately, new generations of gene regulation systems are emerging to solve this problem. For example, the RNA on-switch harnesses an antisense oligonucleotide to regulate ribozyme-base RNA degradation^5^. The X^on^ system utilizes an RNA splicing modulator LMI070 to elicit the inclusion of a start codon-containing exon and protein translation initiation^6^. The pA regulator uses a tetracycline-responsive aptamer to activate gene expression by excluding a synthetic polyA signal in the 5’-untranslated region (5’UTR)^7^. These gene switches are sufficiently small to fit into the limited packaging space of AAV, but whether they could indeed enhance the efficacy and safety of AAV therapy relative to conventional AAV lacks evidence support.

Heart regenerative therapy could potentially benefit from a gene switch because this biological process naturally involves on and off states. Upon cardiac injury, pre-existing cardiomyocytes dedifferentiate into a less mature state, switch on the cell cycle and proliferate to generate new cardiomyocytes. In the following phase when a sufficient number of new cardiomyocytes are produced to repair the injury, the cells turn off the proliferative program, redifferentiate back into the fully mature state to restore heart function^8–10^. A bulk of signal pathways, such as Hippo-YAP^11,12^ and Neuregulin-ErbB^13,14^, are well-established to induce cardiomyocyte dedifferentiation, cell proliferation and heart repair^15–18^. However, prolonged or excessive activation of these pathways cause damage to the heart. For example, transgenic expression of YAP^5SA^, a constitutively active YAP mutant, resulted in cardiomyocyte hyperplasia, thickened ventricle walls and animal death^19^. Prolonged overexpression of wildtype YAP impaired cardiomyocyte mitochondria and caused pathological hypertrophy. Similarly, long-term overexpression of ErbB2 also caused pathological hypertrophy, reduced cardiac output and animal death^20,21^. Interestingly, when the tet-on/off system was utilized to transiently activate ErbB2, the heart showed robust resistance against ischemic injury while the side-effect of prolonged ErbB2 overexpression was circumvented^20,22^. These data strongly indicated that a gene switch was necessary to fulfil safe and effective myocardial regeneration in AAV therapy.

Based on these rationales, this study developed a new gene expression switch responsive to risdiplam, an FDA-approved alternative-splicing modulator that was originally used in the therapy for spinal muscular atrophy^23^. With an inducible dynamic range of over 2000 folds and a regulatable temporal resolution of 2-3 days, this technique was shown to reversibly fine-tune AAV-delivered YAP^5SA^ expression in a cardiomyocyte-specific manner, enhancing myocardial regeneration while greatly reducing YAP^5SA^-associated cardiac toxicity.

## Results

### DreAM, a risdiplam-responsive cassette exon module derived from the murine heart

To determine the endogenous RNA targets of risdiplam in the heart, adult mice were fed with a single dose of 10 mg/kg risdiplam before the heart apexes were collected for RNA sequencing (RNA-Seq, Fig. 1a). Comprehensive alternative splicing hunting^24^ was utilized to determine the risdiplam-elicited splicing events (Supplementary Data 1). After filtering through an array of parameters relevant to an alternative-splicing gene switch, 7 candidate cassette exon modules were selected (Fig. 1a-b, Extended Data Fig. 1a-b) as they harbored robust pseudoexons (PSEs) that could be included in the mature mRNA in a risdiplam-dependent manner.

**Fig. 1:**
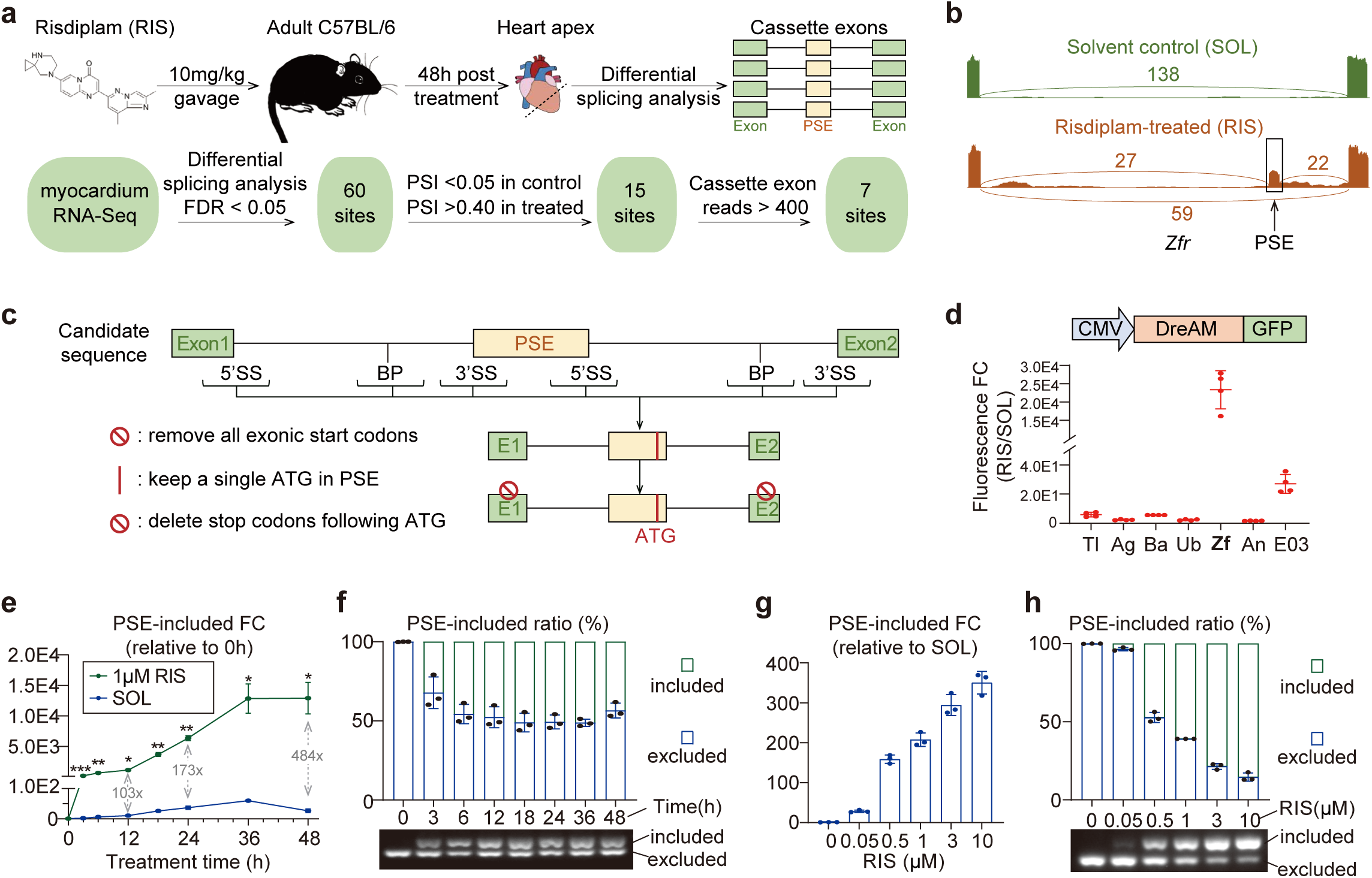
**DreAM is an engineered alternative-splicing module derived from a risdiplam-elicitable cassette exon in murine *Zfr***. **a,** RNA-seq workflow to identify risdiplam-elicitable cassette exons in murine hearts. PSE, pseudoexon. FDR, false discovery rate. PSI, percent spliced-in. n=5 hearts per group. **b,** A representative Sashimi plot showing risdiplam-induced PSE splice-in in *Zfr* intron 22. **c,** Engineering workflow to design a DreAM. The 5’ splice sites (5’SS), 3’ splice sites (3’SS) and branch points (BP) in candidates were predicted according to motif knowledge. Start codons were removed in all exons, adding a single ATG in PSE. All stop codons in frame with the ATG were also ablated. **d,** DreAM candidates were incorporated between a CMV promoter and a GFP reporter. The start codon of GFP was removed, and the ATG in DreAM was aligned in frame with GFP. HEK293T cells were transfected with these reporter plasmids and treated with 1 μM risdiplam (RIS) or an equal amount of solvent (SOL, DMSO) for 24h before baseline and risdiplam-induced GFP signals were quantified. FC, fold change. Tl, Ag, Ba, Ub, Zf, An and E03 stand for DreAM candidates derived from *Tlk2*, *Agrn*, *Bag2*, *Ubap2l*, *Zfr*, *Angel2*, *E030003E18Rik*, respectively. **e,** TaqMan-based RT-qPCR quantification of PSE-included mRNA at different time points during RIS or SOL treatment. Y axis value, FC relative to 0 h; value in grey, FC relative to SOL at the same time points. **f,** RT-PCR quantification (above) and raw images (below) of the ratios of mRNA that include or exclude PSE. **g,** RT-qPCR quantification of PSE-included mRNA after treatment for 24h. **h,** RT-PCR quantification (above) and raw images (below) of the ratios of mRNA that include or exclude PSE after treatment for 24h. **d-h,** mean ± SD, n=3-4 biological repeats.

The alternative-splicing switch was intended to fit into the limited 4.7 kbp packaging space of AAV, thus the key sequences for RNA splicing, including 5’ splice sites (5’SS), branch points (BP) and 3’ splice sites (3’SS), were extracted from candidate modules according to known motif information^25^ and were subsequently assembled into smaller modules less than 1.5 kbp. All exonic start codons were removed, leaving a single start codon in the PSE. This design allowed risdiplam-induced PSE inclusion to control protein translation initiation (Fig. 1c).

The candidate modules were positioned in frame to a start codon-deleted GFP gene to build a reporter driven by the CMV promoter. Plasmids carrying these reporters were transfected into HEK293T cells, which were treated with risdiplam or solvent to examine GFP signals at the on and off states, respectively. The module derived from *Zfr* appeared to exhibit the highest GFP signal fold change between the on and off states (Fig. 1d, Extended Data Fig. 1c-d). We termed this module as the drug-elicitable alternative-splicing module (DreAM), which was used in the following studies.

### Characterization of the DreAM activity in cell culture

A TaqMan probe was designed to specifically target the PSE-exon 2 boundary (Extended Data Fig. 2a) for precise quantification of PSE-included mRNA by real-time quantitative PCR (RT-qPCR). Risdiplam treatment triggered a steady increase of PSE-included mRNA over time in HEK293T cells expressing the DreAM-GFP reporter. Compared to the solvent control, risdiplam elicited over 100-fold increase of the PSE-included mRNA within 3 h treatment. An average of 173- and 484-fold changes could be detected by 24 h and 48 h treatment, respectively (Fig. 1e).

A reverse-transcription PCR (RT-PCR) assay was developed to measure the alternative splicing ratio between the PSE-included and PSE-excluded events (Extended Data Fig. 2b). This assay confirmed that the robust splicing changes could be readily detected after 3 h risdiplam treatment and the PSE-included ratio peaked within 24 h (Fig. 1f).

Serial concentrations of risdiplam were applied to the cells for 24 h before RT-qPCR and RT-PCR were performed to build the dose-effect relationship (Fig. 1g-h). The solubility-limited concentration of 10 μM risdiplam induced ∼351-fold increase relative to solvent control with an PSE splice-in ratio of ∼85% with an estimated EC50 of 0.47 μM.

Next, the impact of DreAM incorporation on gene expression of GFP reporter was assessed. An RT-qPCR assay targeting the GFP coding sequence showed that DreAM significantly reduced total GFP mRNA level as compared to the positive control reporter without DreAM (Extended Data Fig. 2c-d). Consistent with this data, risdiplam maximally elicited GFP fluorescence signal at half of the level expressed by the positive control (Extended Data Fig. 2e-f).

DreAM was validated to regulate GFP reporter when the plasmids were transfected into four cell lines including cat CRFK cells, pig IBRS-2 cells, green monkey VERO cells and mouse NIH3T3 cells (Extended Data Fig. 3). The DreAM-GFP minigene was also packaged into lentivirus and adenovirus, which were validated in six additional cell lines including HEK293T cells, HeLa cells, bull MDBK cells, dog MDCK cells, neonatal rat ventricular cardiomyocytes and human embryonic stem cell-derived cardiomyocytes (Extended Data Fig. 4). Therefore, the DreAM-based gene switch is broadly applicable in various mammalian species, cell types and gene delivery vectors.

### Tunable AAV gene expression via DreAM in vivo

The DreAM-GFP reporter was next packaged into an AAV9 vector. 5×10^10^ vg (vector genome) AAV were subcutaneously injected into postnatal day 1 (P1) mice seven days before a single dose of 10 mg/kg risdiplam was administered intraperitoneally (Fig. 2a). Then the heart, liver and skeletal muscles were collected at each day post risdiplam (DPR) treatment for 5 days (Fig. 2b-d). RT-qPCR showed that the PSE splice-in events were peaked at 1 DPR, which was quickly decayed by ∼80% at 2 DPR. The peak fold changes among the tissues ranked as skeletal muscle>heart>liver. Western blot validated this pulsive kinetics of GFP expression (Fig. 2d). Thus, DreAM could modulate AAV gene expression with a temporal resolution of a couple of days.

**Fig. 2:**
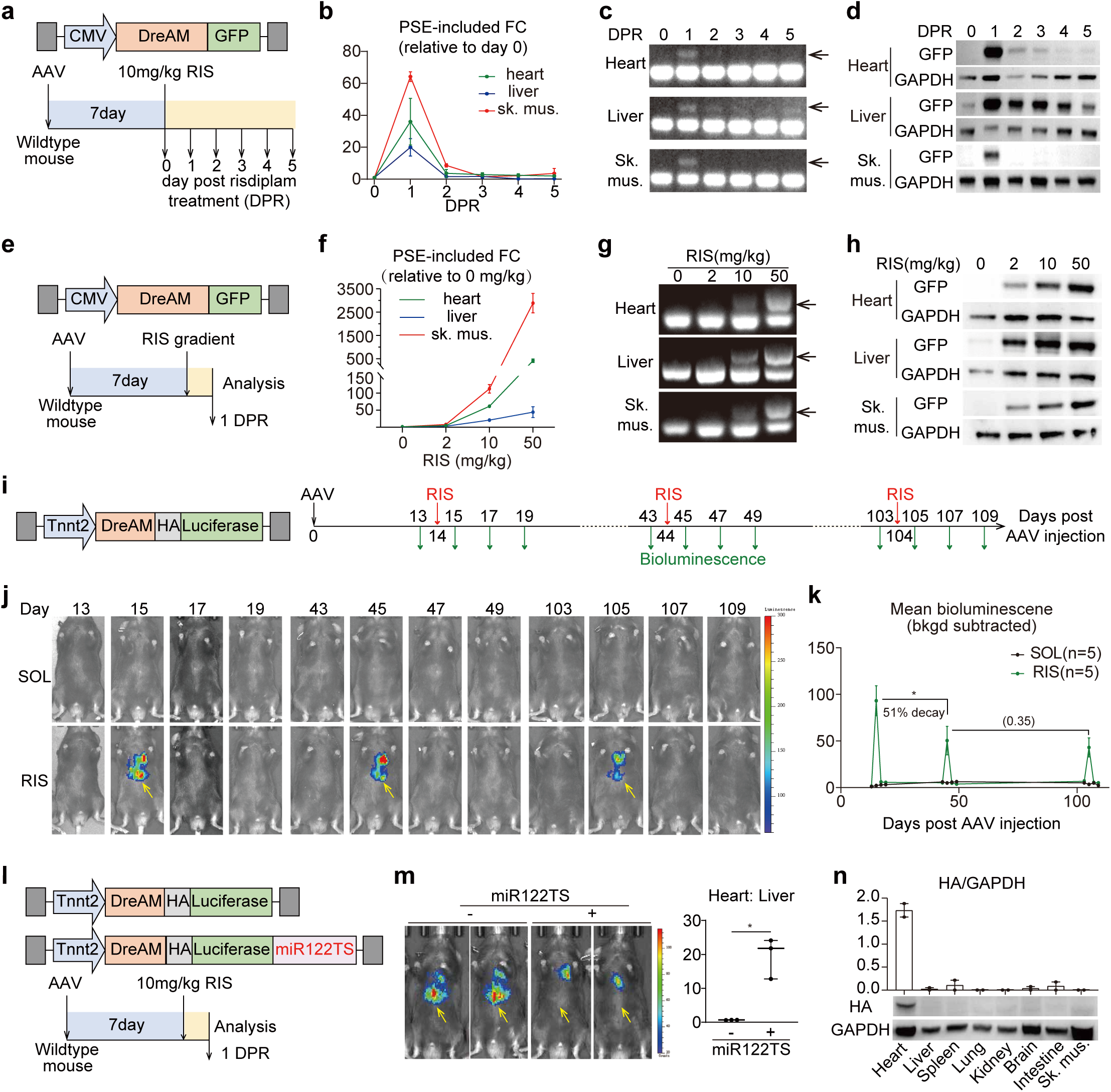
**DreAM dynamically and reversibly switches AAV transgene in response to risdiplam**. **a,** Experimental design to analyze the kinetics of DreAM-induced AAV expression. 5 × 10^10^ vg AAV9 was subcutaneously injected into postnatal day 1 (P1) mice. At P8, a single dose of risdiplam was injected before tissues were collected daily in the following days. **b-c,** RT-qPCR (**b**) and RT-PCR (**c**) analysis of PSE-included mRNA. Sk. mus., skeletal muscle. Arrows point to the PSE-included band. **d,** Western blot quantification of GFP expression at different days after risdiplam treatment. **e,** Experimental design to evaluate the dose-effect relationship of DreAM-regulated AAV expression. 5 × 10^10^ vg AAV9 was subcutaneously injected at P1. Risdiplam was administered at P8, one day before tissues were collected. **f-h,** RT-qPCR (**f**), RT-PCR (**g**) and Western blot (**h**) analysis of AAV transgene expression in response to serial risdiplam dosages. **i,** Experimental design to test the repeatable AAV gene induction by risdiplam. 5 × 10^11^ vg AAV9 was injected into adult mice via tail vein. 10 mg/kg risdiplam was repeatedly fed at 0.5-, 1.5- and 3.5-month post AAV injection. Bioluminescence was detected from same animals before and after each risdiplam treatment. **j-k,** bioluminescence images (**j**) and cardiac signal quantification (**k**). Paired single-tailed student’s t-test, *p<0.05. Non-significant P value in parenthesis. Bkgd, background. Yellow arrows point to the leaky liver signal. **l,** Experimental design to test cardiac specific AAV induction. 5 × 10^11^ vg AAV9 was injected into adult mice via tail vein. miR122TS, microRNA-122 target sequence. **m,** Bioluminescence images and quantification of heart signals relative to liver. Unpaired two-tailed student’s t-test, *p<0.05. n = 3 animals per group. **n,** Western blot images and quantification of HA-Luciferase amounts in various tissues. n = 2 animals per group.

Serial concentrations of risdiplam were applied to examine the dose-effect relationship at 1 DPR (Fig. 2e). At 50 mg/kg, the PSE-included fold change reached over 2500 folds in skeletal muscles and 500 folds in the heart (Fig. 2f-g). Western blot further validated this dose-dependent GFP expression (Fig. 2h). Notably, leaky GFP expression was detected in the liver in the absence of risdiplam, resulting in lower fold changes (Fig. 2b, 2d, 2f and 2h). Thus, DreAM preferentially works in the cardiac and skeletal muscles.

DreAM was next incorporated into a new AAV expressing HA-tagged luciferase driven by the cardiac specific Tnnt2 promoter^26^. Then, 5×10^11^ vg AAV was injected into adult mice via tail veins and risdiplam was administered at 0.5, 1.5 and 3.5 months after AAV injection via gavage feeding (Fig. 2i). Bioluminescence imaging detected luciferase signals at 1 DPR after each risdiplam treatment, which returned to background by 3 DPR. The risdiplam-induced bioluminescence decayed by ∼50% between the first two treatments and reached a steady state thereafter (Fig. 2j-k). Thus, DreAM could be reversibly and repeatedly induced in adult mice via oral administration of risdiplam.

The bioluminescence signals were observed in the liver (Fig. 2j, arrows), confirming the liver leakage of the AAV9-Tnnt2 vectors^27^. To solve this problem, a microRNA-122 targeting sequence (miR122TS) was added to the 3’ untranslated region of the AAV transgene (Fig. 2l)^27^, which robustly eliminated the liver luciferase activity, enhancing the heart-to-liver signal ratio to over 20 folds (Fig. 2m). Western blot further validated the specific AAV expression in the heart but not other organs including liver, spleen, lung, kidney, brain, intestine and skeletal muscle (Fig. 2n). These data demonstrated that DreAM could be combined with cis-regulatory elements to achieve tissue specificity.

### DreAM-based regulation of AAV-delivered YAP activation

In the following studies, DreAM was applied to control YAP^5SA^ expression. When DreAM was directly connected to the YAP^5SA^ gene, risdiplam induced the expression of a protein slightly bigger than the intact YAP^5SA^ protein (Fig. 3a-b). This is because the PSE and exon 2 in DreAM encodes a “junk peptide” at the N-terminus of YAP^5SA^ with unpredictable effects. To avoid this issue, a P2A self-cleaving peptide^28^ was placed between DreAM and YAP^5SA^. P2A cleaved off the junk peptide together with P2A itself, allowing YAP^5SA^ to be expressed with the amino acid sequence almost the same to the intact protein (Fig. 3a-b). 5×10^10^ vg AAV9-DreAM-P2A-YAP^5SA^ vector was injected into P1 mice before daily risdiplam injection was started at P8 (Fig. 3c). Risdiplam-elicited PSE-inclusion and YAP^5SA^ expression in the heart was confirmed by RT-qPCR and Western blot (Fig. 3d-e). No junk peptide was detected in the AAV-delivered YAP^5SA^ protein (Fig. 3e). Immunofluorescence validated the nuclear localization of YAP^5SA^ (Fig. 3f, arrowhead)^29^.

**Fig. 3:**
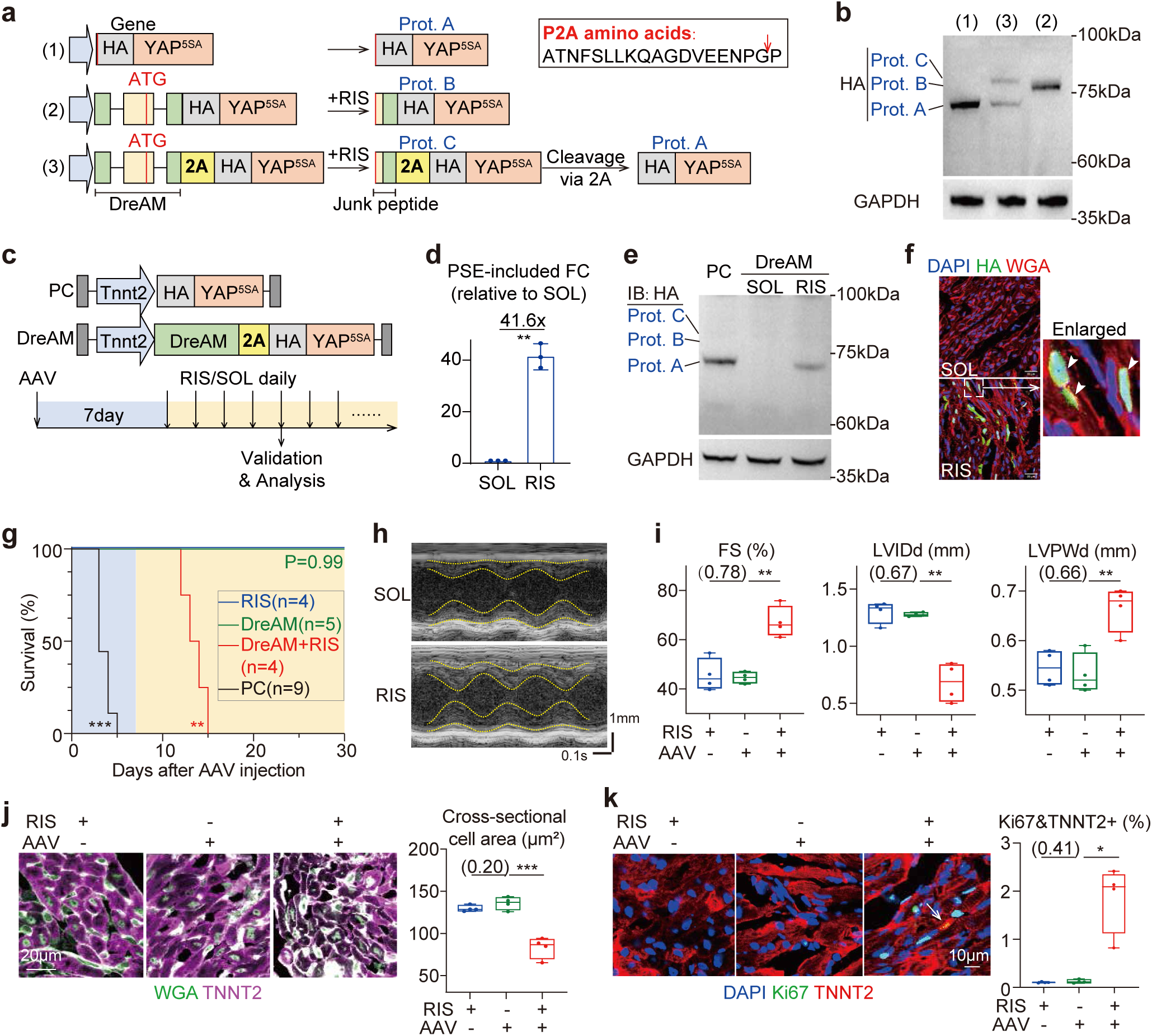
**DreAM can control AAV-delivered YAP^5SA^-triggered myocardial hyperplasia and animal death**. **a,** A diagram showing the design of YAP transgenes and their predicted protein products. (1) Regular HA-YAP^5SA^ gene and protein. (2) DreAM-HA-YAP^5SA^ fusion gene and protein. (3) DreAM-controlled HA-YAP^5SA^ gene and protein cleaved by a 2A peptide. Prot., protein. Prot. A, intact HA-YAP^5SA^ protein. Prot. B, HA-YAP^5SA^ protein with an N-terminal junk peptide coded by DreAM. Prot. C, HA-YAP^5SA^ protein fused to junk peptide via P2A self-cleaving peptide. The P2A amino acid sequence and its cleavage site (red arrow) are indicated in the box. **b,** Western blot analysis of gene expression products by HA-YAP^5SA^ genes in the three formats in in HEK293T cells. **c,** A diagram showing DreAM-YAP^5SA^ and positive control (PC) AAV9 vectors and the experimental design for validation in mice. RIS, 10 mg/kg Risdiplam; SOL, solvent control. 5 × 10^10^ vg AAV was subcutaneously injected at P1 and RIS/SOL were intraperitoneally injected at P8. **d,** TaqMan RT-qPCR quantification of PSE-included transcripts in the heart. Mean ± SD. n = 3 animals per group. **e,** Western blot analysis of HA-YAP^5SA^ expression in the heart. **f,** Immunofluorescence analysis of cardiac cryosections. Arrowhead, HA-positive nuclei. Scale bar, 20 μm. **g,** Survival analysis of animals treated as shown in (**c**). Log-rank test compared to the RIS control group, **p<0.01, ***p<0.001. **h-i,** Echocardiography (**h**) and analysis of heart structure and function (**i**). FS, fractional shortening. LVIDd, left ventricular inner diameter end diastole. LVPWd, left ventricular posterior wall thickness end diastole. **j-k,** Immunostaining of cardiac cryosections and the analysis of cardiomyocyte cross-section area (**j**) and Ki67 (**k**). WGA, wheat germ agglutinin. In **k**, the arrow points to a Ki67+; TNNT2+ cell. In **d, i, j, k,** unpaired two-tailed student’s t-test, *p<0.05, **p<0.01, ***p<0.001. Non-significant P value in parenthesis. n = 3-4 animals per group.

In mice treated with a positive control AAV-YAP^5SA^ vector in the absence of DreAM, all mice died by day 5 after AAV injection (Fig. 3g), confirming the lethal toxicity of uncontrolled YAP^5SA^ expression^19^. By contrast, DreAM-controlled AAV-YAP^5SA^ did not cause animal death unless risdiplam induction lasted for more than 4 days (Fig. 3g). Echocardiogram showed increased cardiac fractional shortening (FS), decreased end diastolic left ventricular inner diameter (LVID) and increased end diastolic left ventricular posterior wall thickness (LVPWd) when risdiplam continuously turned on YAP^5SA^ (Fig. 3h-i). In this group, wheat germ agglutinin (WGA) staining revealed the reduced cardiomyocyte cross-sectional area (Fig. 3j) and Ki67, a cell proliferation marker, was significantly increased in TNNT2-positive cardiomyocytes (Fig. 3k). Therefore, daily risdiplam treatment could switch on DreAM and induce continuous YAP^5SA^ activation, myocardial hyperplasia and animal death, similar to the published observation in transgenic mice^19^.

### DreAM-regulated YAP^5SA^ expression to build a dedifferentiation-redifferentiation cycle

In the following studies, risdiplam treatment was reduced to two days to achieve transient AAV-YAP^5SA^ expression. Western blot detected peak YAP^5SA^ expression at 1 DPR, which quickly returned to baseline by 11 DPR (Fig. 4a). These two timepoints were named as the ON and OFF stages in the following analyses. RNA-Seq of heart tissues showed that YAP^5SA^ altered the expression of 1627 genes at the ON stage, but the number and fold changes of differentially expressed genes (DEGs) were greatly reduced at the OFF stage (Fig. 4b, Supplementary Data 2-3). This stage-dependent effect was confirmed by the principal component analyses (Extended Data Fig. 5a). Gene set enrichment analysis (GSEA)^30^ confirmed the upregulation of a commonly used YAP signaling gene set^31^ at the ON stage, which became less prominent at the OFF stage (Fig. 4c).

**Fig. 4:**
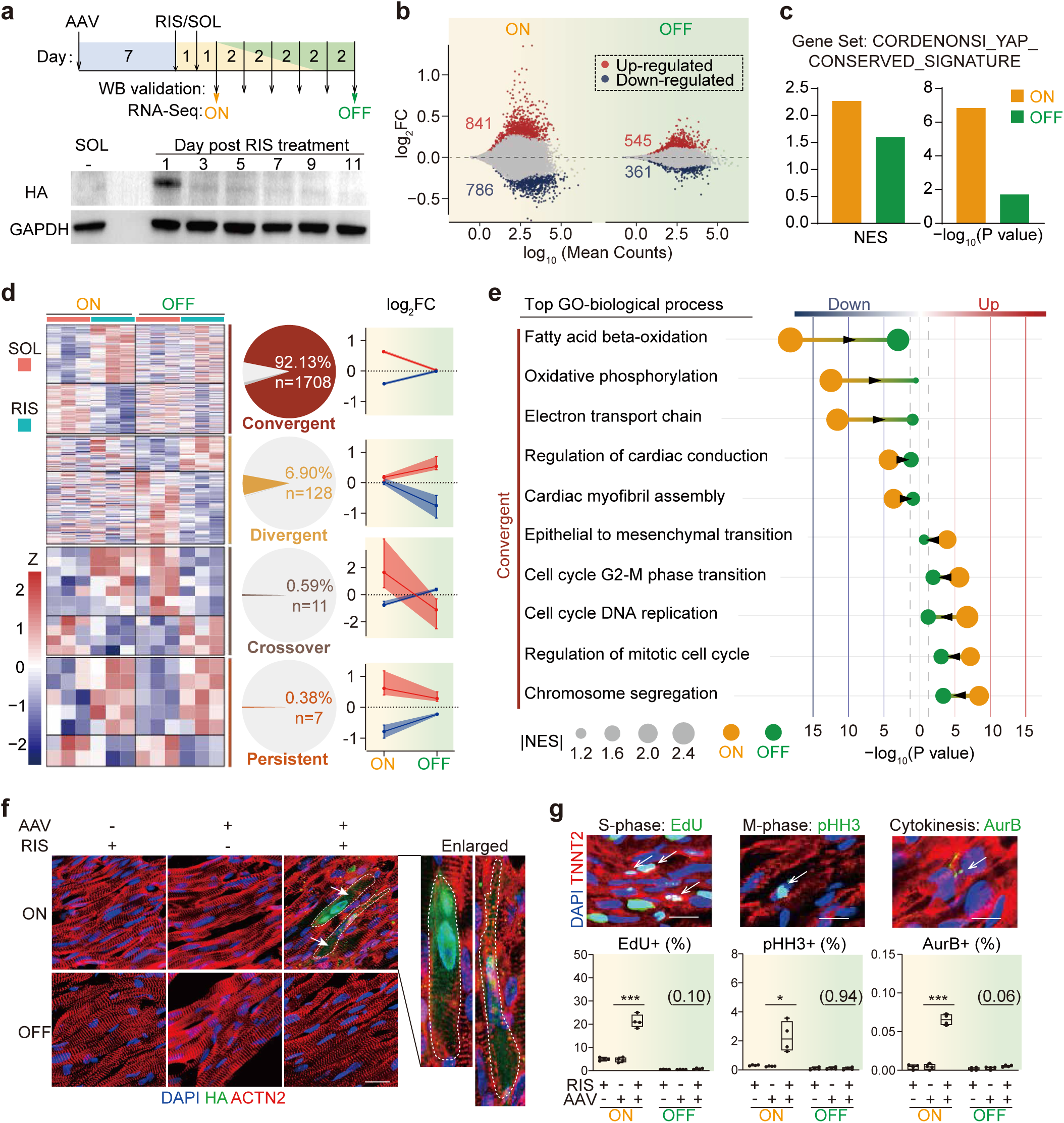
**DreAM transiently activates AAV-YAP^5SA^ and builds a dedifferentiation-redifferentiation cycle**. **a,** Experimental design to explore the effect of DreAM-regulated transient YAP^5SA^ expression. Western blot below the diagram shows HA-YAP expression at serial timepoints post risdiplam treatment. **b,** MA-plot of RNA sequencing data between RIS and SOL at the ON and OFF timepoints. DESeq2-generated adjusted p value < 0.05 as cutoff for statistical significance. **c,** GSEA analysis using the cordenonsi_yap_conserved_signature gene set from MSigDB. NES, normalized enrichment score. **d,** Heatmap and expression profiles of differentially expressed (DE) genes at the two timepoints. In heatmaps, rows represent genes, and columns represent biological repeats. Dynamic expression patterns are divided into 4 groups: convergent (DE at the ON stage, not DE at the OFF stage), divergent (not DE at the ON stage, DE at the OFF stage), crossover (DE at the ON and OFF stages in opposite directions) and persist (DE at the ON and OFF stages in the same direction). Pie charts indicate gene number and ratio in each group. Line charts show the gene log_2_FC changes in each group. Median ± 95% confidence interval in line charts with up- and down-regulated genes separately analyzed. FC, fold change. **e,** GSEA analysis of convergent gene ontology (GO) biological process gene sets at ON and OFF timepoints. Arrowheads point to the directions of changes over time. **f,** Representative immunofluorescence images of sarcomere Z-line marker ACTN2. Dashed lines delineate cell boundaries. Arrows point to HA-positive cells. Scale bars, 20Cμm. **g,** Representative images and quantification of cardiomyocyte cell cycle markers. Arrows point to cell cycle markers in TNNT2-positive cells. Scale bar, 10Cμm. Student’s t-test, *p<0.05, ***p<0.001. Non-significant P value in parenthesis. N = 4-5 animals per group.

Among the dynamically changed DEGs, 92.1% were significantly changed at the ON stage and reversed at the OFF stage (Fig. 4d). GSEA of signature gene sets for immature and mature cardiomyocytes^32^ showed that the immature markers, such as *Myh7*, were up-regulated while the mature markers, such as *Myh6*, were down-regulated at the ON stage. Both the mature and immature gene changes returned to baseline at the OFF stage (Extended Data Fig. 5b-c). Gene ontology (GO) analyses showed that fatty acid metabolism, oxidative phosphorylation and myofibril assembly were the major biological processes that were temporarily down-regulated at the ON stage, while terms relevant to cell cycle were temporarily up-regulated (Fig. 4e, Supplementary Data 4-5). These findings were further validated using HALLMARK^33^ and KEGG^34^ gene sets (Extended Data Fig. 5d). Although these key dedifferentiation gene sets showed clear reversion at the OFF stage, a small but statistically significant P value could be detected. This result implied the presence of residual dedifferentiation that was resistant to reversal, agreeing to a recent study^22^. Risdiplam alone did not cause these gene expression changes (Extended Data Fig. 5e, Supplementary Data 6-7), ruling out the influence by risdiplam itself.

As phenotypic validations of the gene expression change, immunostaining of ACTN2, a marker of sarcomere Z-line^32^, was performed on heart tissues. At the ON stage, sarcomere disassembly was observed in cardiomyocytes with detectable HA-YAP^5SA^ expression. However, no disruption of sarcomere organization could be found at the OFF stage (Fig. 4f). EdU incorporation, histone H3 phosphorylation and the midbody localization of Aurora B were next examined in TNNT2-positive cardiomyocytes to identify cells entering the S-phase, mitosis and cytokinesis, respectively. These assays showed transient increase of cardiomyocyte proliferation at the ON stage, but not at the OFF stage (Fig. 4g). Overall, these data demonstrated DreAM as a powerful technique to enable transient AAV-YAP^5SA^ expression for only a couple of days, which was sufficient to build a full cycle of cardiomyocyte dedifferentiation, proliferation and redifferentiation.

### DreAM-based regeneration therapy for myocardial infarction in mice

The AAV-DreAM-YAP^5SA^ vector was next tested in the therapy for myocardial infarction (MI) in mice. MI was induced by permanent ligation of the left anterior descending coronary artery, which was validated by acutely decreased ejection fraction (EF) at day 2 post MI by echocardiogram. Daily risdiplam (10 mg/kg) treatment for a month in the MI model did not alter cardiac dysfunction as compared to solvent control (Extended Data Fig. 6a). Masson trichrome staining detected no effect of risdiplam on MI-induced fibrotic area (Extended Data Fig. 6b). Blood biomarkers for liver and muscle damages and lipid metabolism were not altered by risdiplam (Extended Data Fig. 6c-d). Hematoxylin-eosin staining detected no histological changes in liver, spleen, lung, kidney, brain and skeletal muscle (Extended Data Fig. 6e). Thus risdiplam alone did not perturb the progression of MI-induced cardiac pathogenesis.

Next, 5×10^11^vg AAV was administered to MI mice via tail veins at day 3 post MI. Five days later, risdiplam or solvent was fed for two days before heart tissues were collected for immunofluorescence validation (Fig. 5a). In hearts treated with the AAV-YAP^5SA^ positive control vector or the AAV-DreAM-YAP^5SA^ vector induced by risdiplam, HA-YAP^5SA^ signal was predominantly detected at the border zone between the infarcted region and the non-infarcted myocardium (Fig. 5b and Extended Data Fig. 7a). No HA signal could be observed in the negative controls (Extended Data Fig. 7a). EdU incorporation assay also validated the induced cell proliferation preferentially at the border zone (Fig. 5c and Extended Data Fig. 7a).

**Fig. 5:**
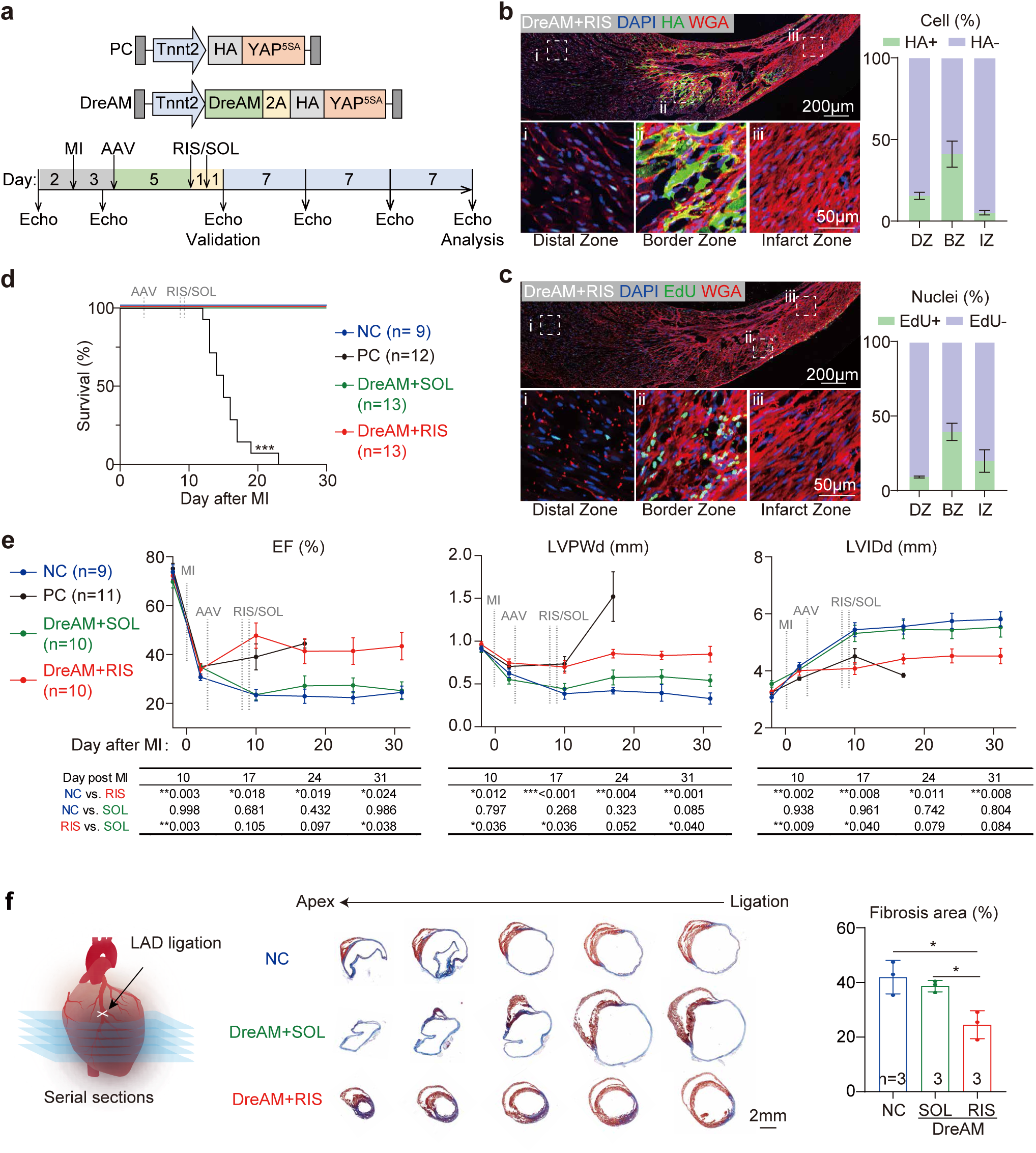
**DreAM-controlled YAP^5SA^ expression ameliorate myocardial infarction-induced myocardial injury while circumventing YAP^5SA^-induced animal death**. **a,** Experimental design to explore the effect of DreAM-regulated transient YAP^5SA^ expression on myocardial infarction in mice. **b-c,** Fluorescence images of heart sections. MI mice were treated with 5×10^11^vg AAV-DreAM-YAP^5SA^ via tail vein at day 3 post infarction. At day 8 after MI, the mice were treated with risdiplam and EdU on two consecutive days before the hearts were collected for validation at day 10 post MI. DZ, distal zone; BZ, border zone; IZ, infarct zone. **d,** Survival curve of MI mice. Log-rank test compared to the NC group, ***p<0.001. NC, negative control. PC, positive control. **e,** Serial echocardiogram analysis and pair-wise statistical analysis via mixed-effect ANOVA with Tukey’s multiple comparison correction. Adjusted P value was shown in the table, *p<0.05, **p<0.01, ***p<0.001. **f,** Masson’s trichrome staining on serial cardiac sections of the MI hearts and quantification of fibrosis area. Student’s t-test, *p<0.05.

All animals that were treated with the AAV-YAP^5SA^ positive control vector died by 3 weeks after MI (Fig. 5d), even though their cardiac EFs were significantly increased (Fig. 5e). Their hearts exhibited massively increased left ventricular posterior wall thickness (LVPW) and decreased left ventricular inner diameter (LVID) beyond the normal values (Fig. 5e). By contrast, in AAV-DreAM-YAP^5SA^-treated mice transiently induced by risdiplam, no animals died and cardiac EFs were significantly restored. LVPW and LVID were returned to a relatively normal value (Fig. 5e). The fibrotic scar size was also reduced in this group (Fig. 5f). One month after MI, no significant changes of blood biomarkers for liver and muscle damages could be detected in the therapeutic group (Extended Data Fig. 7b). No histological changes in the liver could be observed (Extended

Data Fig. 7c). Together, these data showed that DreAM enabled tunable AAV- YAP^5SA^ expression that could achieve effective heart regeneration and recovery of heart functions while significantly reducing the risks associated with uncontrolled and prolonged expression.

## Discussion

An ideal gene therapy requires the therapeutic gene to be expressed at the correct time and level. However, current AAV therapy lacks such a component to fine-tune gene expression. To solve this problem, this study repurposed the FDA-approved RNA splicing inducer, risdiplam, to build the DreAM gene switch (Fig. 6a). DreAM was validated in 10 cell types from 8 mammalian species using 4 forms of vectors. When delivered by AAV, DreAM controlled gene expression with an inducible change up to 2000 folds, depending on risdiplam dosage and the targeted organ. DreAM is compatible with regular cis-regulatory elements for tissue-specific regulation. A single dose of risdiplam could elicit peak gene expression within a day, which quickly diminished in a couple days, creating high versatility in manipulating gene expression timing.

**Fig. 6:**
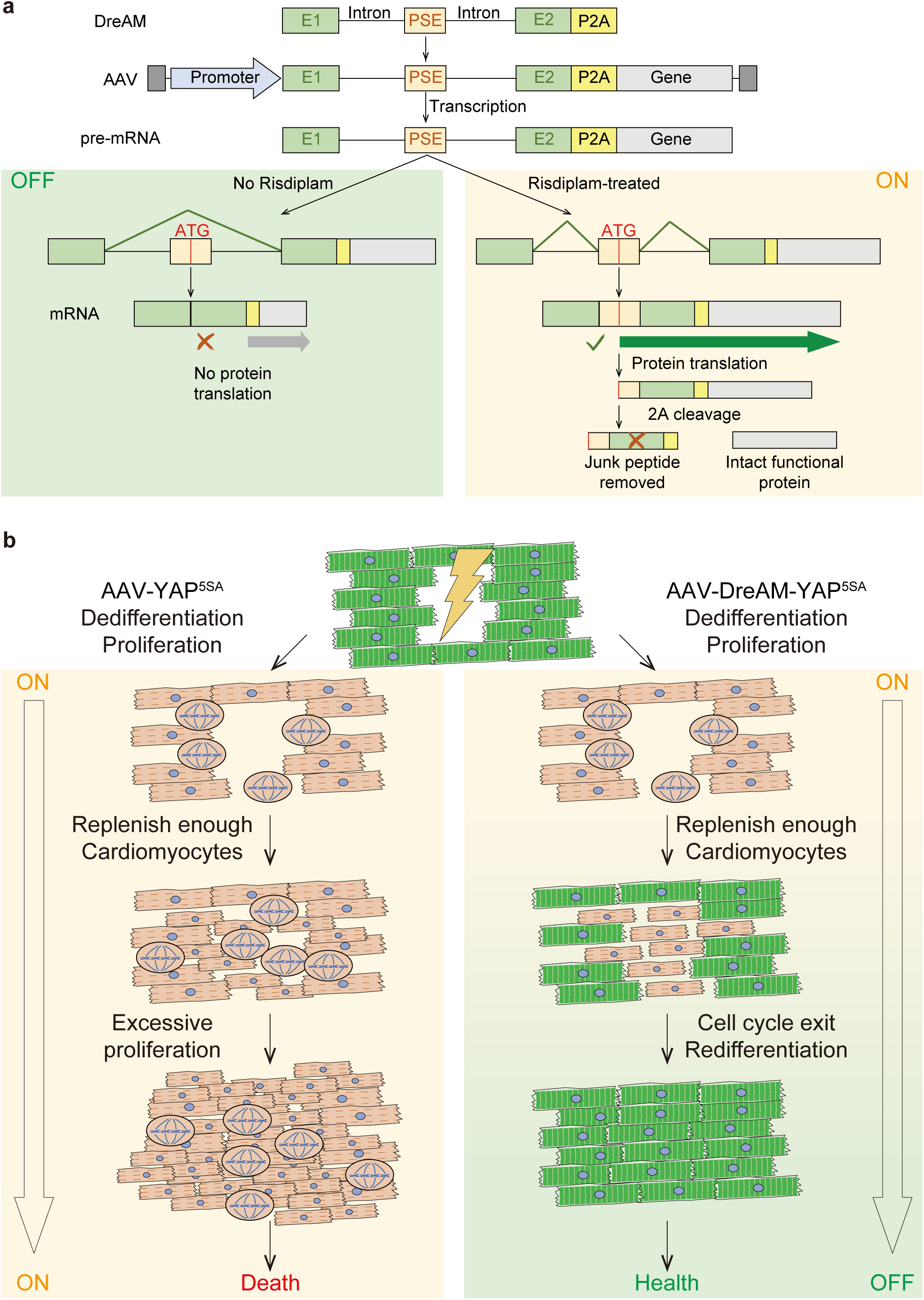
**A graphical summary**. **a,** The working mechanism of DreAM. In the absence of risdiplam, the splicing of DreAM results in the formation of direct E1-E2 junction, excluding PSE. Due to the lack of start codon, mRNA in this form does not translate into protein. By contrast, with the presence of risdiplam, PSE was spliced into mature mRNA. The start codon in PSE initiated protein translation. A P2A self-cleavage peptide was positioned between DreAM and the transgene in frame, so the DreAM and P2A peptides were removed after translation, producing an almost intact protein of interest. **b,** The rationale of DreAM-realized myocardial regeneration therapy. In cardiomyocytes transduced by conventional AAV-YAP^5SA^ vectors, YAP^5SA^ expression lasts for a long time, resulting in persistent cardiomyocyte proliferation, myocardial hyperplasia followed by animal death. By contrast, DreAM could control the expression time and level of AAV- YAP^5SA^. After transient YAP^5SA^ expression, cardiomyocytes were allowed to redifferentiate back to the mature and functional state to restore heart functions in the MI model.

Several AAV-compatible gene switches have been recently reported^5–7^, but little evidence was available to prove that these gene switches could practically enhance therapeutic efficacy and safety as compared to conventional AAV vectors. In this study, DreAM-mediated cardiac regeneration therapy provided the key proof of concept that AAV gene therapy could indeed tremendously benefit from a gene switch. Natural heart regeneration is biphasic, including the ON state of cardiomyocyte dedifferentiation and proliferation being followed by the OFF state of cardiomyocyte redifferentiation and maturation. DreAM realized transient AAV-YAP^5SA^ expression and opened the door to fully mimic the dedifferentiation-redifferentiation cycle in a therapeutic platform, which circumvented the lethal side-effects associated with prolonged YAP^5SA^ expression delivered by conventional AAV (Fig. 6b). Thus, DreAM created an unprecedented opportunity to improve cardiac gene therapy suffering from uncontrolled AAV expression^35^.

Although DreAM successfully realized AAV-YAP^5SA^-based cardiac repair, further improvements are necessary in the future studies to enhance the outcome of this regenerative therapy. First of all, because continuous AAV-DreAM-YAP^5SA^ induction by daily risdiplam treatment caused animal death in four days, AAV-DreAM-YAP^5SA^ expression was limited to only 2-3 days in the therapeutic study. This time window was too short to trigger more extensive regeneration and better restoration of heart function. Potential solutions included the use of a milder YAP mutant such as YAP^S127A^ ^11^ in combination with more extensive titration of the AAV dosage and risdiplam administration.

The FDA-approved daily dosage of risdiplam for SMA treatment in human was up to 0.25 mg/kg^36^, which could be converted to ∼3 mg/kg in mice via a standard body weight/body surface area conversion^37^. This calculated value is lower than the 10 mg/kg dosage used in our therapeutic study, questioning if the current FDA-accepted risdiplam dosage was sufficient for clinical applications of the DreAM platform. Unlike the SMA therapy that involves years of risdiplam treatment, DreAM requires risdiplam treatment for only several days or weeks to achieve pulsive AAV expression. Thus, a new safety study would be necessary to determine if risdiplam could be applied at a higher dosage than the current FDA guideline while the duration of risdiplam treatment was reduced. In addition, new DreAMs that are more sensitive to risdiplam than the current DreAM could be engineered in the future so less risdiplam is used in gene therapy.

## Methods

### Risdiplam

For mouse experiments, 10 mg risdiplam (T16757, TargetMol) powder was resuspended in 1 ml DMSO (196055, MPbio) before 5-minute sonication was applied. Then, 4 ml PEG300 (GR1216, HarveyBio) was added to the mixture, which were swirled on a Vortex mixer until the power was fully dissolved. Next, 0.5 ml TWEEN-80 (T118633, Aladdin) and 4.5 ml saline (HDLS001119, Solarbio) were added sequentially to generate the solution (1 mg/ml) for animal treatment.

For cell culture experiments, risdiplam powder was dissolved in DMSO (196055, MPbio) to produce the stock solution with a final concentration of 10 mM. All the stock and working solutions were aliquoted and stored in -80°C refrigerator for no longer than 1 month.

### Animals

Animal experiments were in conformity to the mouse protocol authorized by the IACUC of Peking University with the approval number DLASBD0203. C57BL/6 mice were obtained from the Department of Laboratory Animal Science of Peking University Health Science Center, and were kept in ventilated cages with appropriate temperature (23 ± 1°C), suitable humidity (50 ± 5%), controlled illumination (12h dark/light cycle), and unrestricted access to water and food.

The administration route of risdiplam and AAV was determined by mouse ages. Intraperitoneal injection was performed on mice younger than 2 weeks, while gavage feeding was conducted for older mice. AAV was injected to P1 mice subcutaneously. Tail vein injection was performed on adult mice for AAV delivery.

Anesthesia was performed via 3% isoflurane (R510-22-10, RWD) inhalation. For mice younger than 7 days, animals were sacrificed by decapitation. Older animals were euthanized by cervical dislocation following anesthesia.

### RNA-seq and data analysis

Total RNA was purified using TransZol Up Plus RNA Kit (ER501-01-V2, TransGene) with genome DNA removed. RNA-Seq was then performed by the Geekgene or GenePlus in Beijing, China. RNA purification and purity were evaluated by Qubit 4. RNA integrity was assessed using Agilent 2100 Bioanalyzer. Libraries were constructed by the Hieff NGS Ultima Dual-mode mRNA Library Prep Kit for Illumina (12301, Yeasen), and sequenced on the Illumina NovaSeq 6000 Platform or DNBSEQ-T7 platform. An average of 27M reads per library was acquired with the 2× 150bp paired-end setup. FastQC (v0.11.9) provided the quality control checks of the raw reads. After trim_galore (v0.6.7) was employed, the trimmed reads were aligned versus mouse genome version GRCm38/mm10 using STAR (v2.7.9a)^38^.

For differential splicing analysis, Samtools (v1.13)^39^ was used to generate duplicate-removed, indexed BAM files. Alternative splicing was analysis by CASH (v2.2.1)^24^. To narrow down high-quality sites for further engineering of DreAMs, the differential splicing events were filtered by adjusted P value <0.05, PSI (percent-spliced-in) < 0.05 in the control group, PSI > 0.40 in the risdiplam-treated group and total cassette exon reads > 400. Sashimi plots were generated by IGV (v2.16.2)^40^.

For differential expression analysis, read counts were calculated by featureCounts (v2.0.1)^41^. Differentially expressed genes were identified utilizing DESeq2 (v1.42.0)^42^. Adjusted P value less than 0.05 was considered statistically significant. Gene lists pre-ranked by [-sign(FC)·lg(P value)] were generated to perform GSEA using fGSEA (v1.28.0)^43^.

### Plasmids

Plasmid names and full sequences are summarized in Supplementary Table 1. Plasmids for cell culture transfection and AAV packaging share the same backbone. DNA sequences for luciferase (addgene#69915)^11^, miR122TS (addgene#117384)^27^ and YAP^5SA^ (addgene#33093)^44^ were synthesized by GENEWIZ. The Tnnt2 promoter of AAV-Tnnt2-GFP-v2 (addgene#165036)^32^ was replaced by the CMV promoter to produce an AAV-CMV-GFP plasmid. Then a multicloning site (MCS) was synthesized and inserted into AAV-CMV-GFP though NcoI sites to construct the AAV-CMV-MCS-(no ATG)-GFP negative control plasmid. The engineered DreAM candidates of Tlk2, Agrn, Bag2, Ubap2l, Zfr, Angel2 and E030003E18Rik were synthesized and inserted into the SacII and EcoRI sites of AAV-CMV-MCS-(no ATG)-GFP to generate the reporter plasmids. The CMV promoter of AAV-CMV-DreAM-GFP was replaced by the human Tnnt2 promoter to produce AAV-hTnnt2-DreAM-GFP. The GFP coding sequence was next replaced by synthesized P2A-HA-luciferase, P2A-HA-luciferase-3×miR122TS, HA-YAP^5SA^ and P2A-HA-YAP^5SA^ sequences to build the corresponding plasmids.

For lentivirus plasmids construction, the 3×Flag-MCS fragments in LV3-SFFV-3×Flag-MCS-GFP (QZ35395, QIAGEN) was replaced by DreAM though NotI and SpeI sites to produce the LV3-SFFV-DreAM-GFP plasmid. For adenovirus production, AdV-CMV-DreAM-GFP was generated using the the AdEasy adenoviral system (Agilent) by Hanbio in Shanghai, China.

### Cell culture and fluorescence analysis

HEK293T, CRFK, IBRS-2, VERO, NIH3T3, HeLa, MDBK and MDCK cells were obtained from Procell, Wuhan, China. See Supplemental Table 2 for more information. Their species were validated by RNA-seq in the lab before experiments were performed. These cells were cultured in high glucose DMEM (HK2109.07, HUANKE) containing 10% FBS (SE100-011, VISTECH) and 1×penicillin-streptomycin solution (BC-CE-007, Bio-channel) at 37°C and 5% CO_2_ with saturating humidity. Cells were plated at a density of 1.5×10^5^ cells per well in 24-well plates 12 hours before plasmid transfection or virus transduction.

To isolate neonatal rat ventricular myocytes (NRVM), the ventricles of neonatal rat hearts were minced in HBSS (PB190321, Procell) and enzymatically dissociated in lysis buffer containing 0.1% trypsin (T8150, Solarbio) and 0.056% collagenase (LS004177, Worthington Biochemical). The cells were filtered through a 40 μm strainer (93040, SPL life sciences) and were then plated for 1.5 hours. Unattached cells were resuspended with fresh medium (high glucose DMEM containing 10% FBS, 1×penicillin-streptomycin and BrdU) and plated at a density of 1×10^6^ cells/ml. BrdU was removed from medium before adenovirus transduction.

Human embryonic stem cells (hESCs) (H7, WAe007-A, NIHhESC-10-0061) were cultured on Matrigel (Corning, 354277) in stem cell maintenance medium (Cellapy, CA1014500) and were passaged with DissoEasyII (Cellapy, CA1023100). ESCs were differentiated into cardiomyocytes using a classic two-small molecule protocol^45,46^. In detail, when PSCs reach 70% confluency, medium was changed to RPMI 1640 medium (Gibco, 61870) with B27-insulin (Gibco, A18956-01) and 5 μM CHIR99021 (StemCell Technologies, 72054). After 48h, medium was replaced by RPMI+B27 medium adding 5 μM IWR-1-endo (StemCell Technologies, 72564). At day 4 of the differentiation, hESC-CMs were cultured in RPMI+B27 medium. hESC-CMs cultured for 30 days were used for experiments.

Lipo8000 (C0533, Beyotime) or jetPRIME (101000046, Polyplus) was used following manufacturer’s instructions during plasmid transfection. Virus transduction was performed with 0.1 % polybrene (40804ES76, Yeasen). See Supplementary Table 2 for more information. After 6 hours of plasmid transfection or 24 hours of virus transduction, cells were treated with fresh medium containing the inducers or solvent.

The fluorescence intensity of cultured cells was measured on a CLARIOstar Plus multi-mode microplate reader (BMG LABTEC) at room temperature. Alternatively, images were taken under a fluorescence microscope (BZ-X810, KEYENCE) before the images were quantified by ImageJ (version 1.54f).

### PCR-based gene expression analysis

Total RNA was purified using TransZol Up Plus RNA Kit (ER501-01-V2, TransGene) with genome DNA removed. RNA was reverse-transcribed with HiScript III All-in-one RT SuperMix (R333-01, Vazyme Biotech). RT-PCR was performed using 2×Taq PCR Mix (TIANGEN Biotech) and separated by agarose gel electrophoresis.

For classic SYBR qPCR, 3 technical replicates were assayed using the AriaMx Real-Time PCR System (Agilent Technologies) and 2×Taq Pro Universal SYBR qPCR Master Mix (Q712-02, Vazyme Biotech). TaqMan-based qPCR was performed using QuantStudio 3 Real-Time PCR System (Thermo Fisher Scientific) and AceQ qPCR Probe Master Mix (Q112-02, Vazyme Biotech). TaqMan probes were synthesized by GenScript Biotech. Relative quantitation of gene expression was performed by normalizing to *Gapdh*. Primers and probes are listed in Supplementary Table 3.

### Western blot

Cells or tissues were washed by ice-cold PBS prior to lysis in RIPA Buffer (P0012B, Beyotime Biotechnology). After centrifugation, supernatant was diluted in 4× SDS-PAGE loading buffer (B1016, Solarbio). After being boiled at 70°C for 10 minutes, proteins were separated on 4–12% SurePAGE gradient gel (M00725, GenScript), transferred to Immobilon-P PVDF membrane (IPVH00010, Merck Life Science), and blocked by 5% Non-Fat milk/TBST (Tris buffered saline with 0.1 % Tween 20) at 25°C for 1 hour.

Membranes were incubated with the primary antibodies overnight at 4 °C, followed by 3 × 5 minutes TBST washes. The membranes were probed by horse radish peroxidase (HRP)-conjugated secondary antibodies for 1 hour at room temperature, followed by 3 × 5 minutes TBST washes. After adding ECL Western Blotting Substrate (PE0010, Solarbio), chemiluminescence were detected by an iBright CL1500 Imaging System (Thermo Fisher Scientific) or a Universal Hood II Molecular Imager GEL System (Bio-Rad Laboratories). Antibodies used in this study are listed in Supplementary Table 4. All uncropped western blots can be found in source data.

### Immunofluorescence and imaging analysis

For immunostaining, tissue sections were incubated with primary antibodies diluted in blocking buffer overnight at 4 °C. After the primary antibodies were washed off, sections were incubated with secondary antibodies and dyes overnight at 4 °C. The cells were next washed with PBS and mounted with ProLong Diamond Antifade Mountant (P16961, Thermo Fisher Scientific) before imaging. All antibodies and dyes are listed in Supplementary Table 4.

For the EdU assay, mice were injected intraperitoneally with 50 μg/g EdU in PBS 48 and 24Chours before sample collection. EdU-positive cells were detected using the Yefluor 488 EdU Imaging Kit (40275ES76, Yeasen) according to the manufacturer’s protocol.

Confocal fluorescence images were taken using an Olympus FV3000 confocal laser scanning microscope equipped with a ×40/1.406 silicone-oil objective. Cell size and number and fluorescence intensity were measured using the ImageJ (version 1.54f).

### Myocardial infarction

Male C57BL/6 mice of 8∼12-month ages were used in myocardial infarction. Animals were anesthetized via 3% isoflurane inhalation and intubated with 1.5% isoflurane for sustained anesthesia and ventilation (ALC-ANE6, ALCBIO). After thoracotomy on a warm pad, the left anterior descending coronary artery was ligated with a 7-0 suture. Infarction was confirmed by a pale color in the apical area. After the ligation, the chest was closed and the animals were allowed to recover on a heating pad for 20 minutes. Echocardiogram was performed two days before and after MI surgery. An EF < 45% cutoff was set to exclude mice with failed or weak MI. Then, the mice were randomly grouped for following procedures. Deaths occurred within the first three days after MI were considered secondary to the surgery and excluded in survival analysis.

## Statistical analysis

Statistical analysis and plotting were performed using GraphPad Prism version 9.5.1 for windows, Massachusetts USA, www.graphpad.com. Normal distribution was tested via the Shapiro–Wilk test. In principle, Comparisons were performed using Student’s t-test for normally distributed data. Kaplan-Meier curves with log-rank test were used to evaluate survival. Tukey’s methods were used for multiple comparison corrections. See more details in figure legends for statistical analysis methods. The following values were considered to be statistically significant:

*P<0.05, **P<0.01, ***P<0.001. Unless otherwise stated, dependent on the data distribution, data were presented in figures using the mean ± standard deviation (SD) in bar plots or median with 25th and 75th percentiles (Q1-Q3) and maximal/minimal values (whiskers) in box plots. Dots to represent raw data points.

## Data availability

Plasmids are available at Addgene. Next-generation sequencing data are available at GEO (GSE270445). Other data supporting the findings in this study are included in the main article and source data.

## Code availability

All software used in this study was from public software packages described in Methods and detailed in the accompanying references.

## Ethics declarations

## Competing interests

Patents are filed relating to the data presented.

---

**Extended Data Fig. 1:**
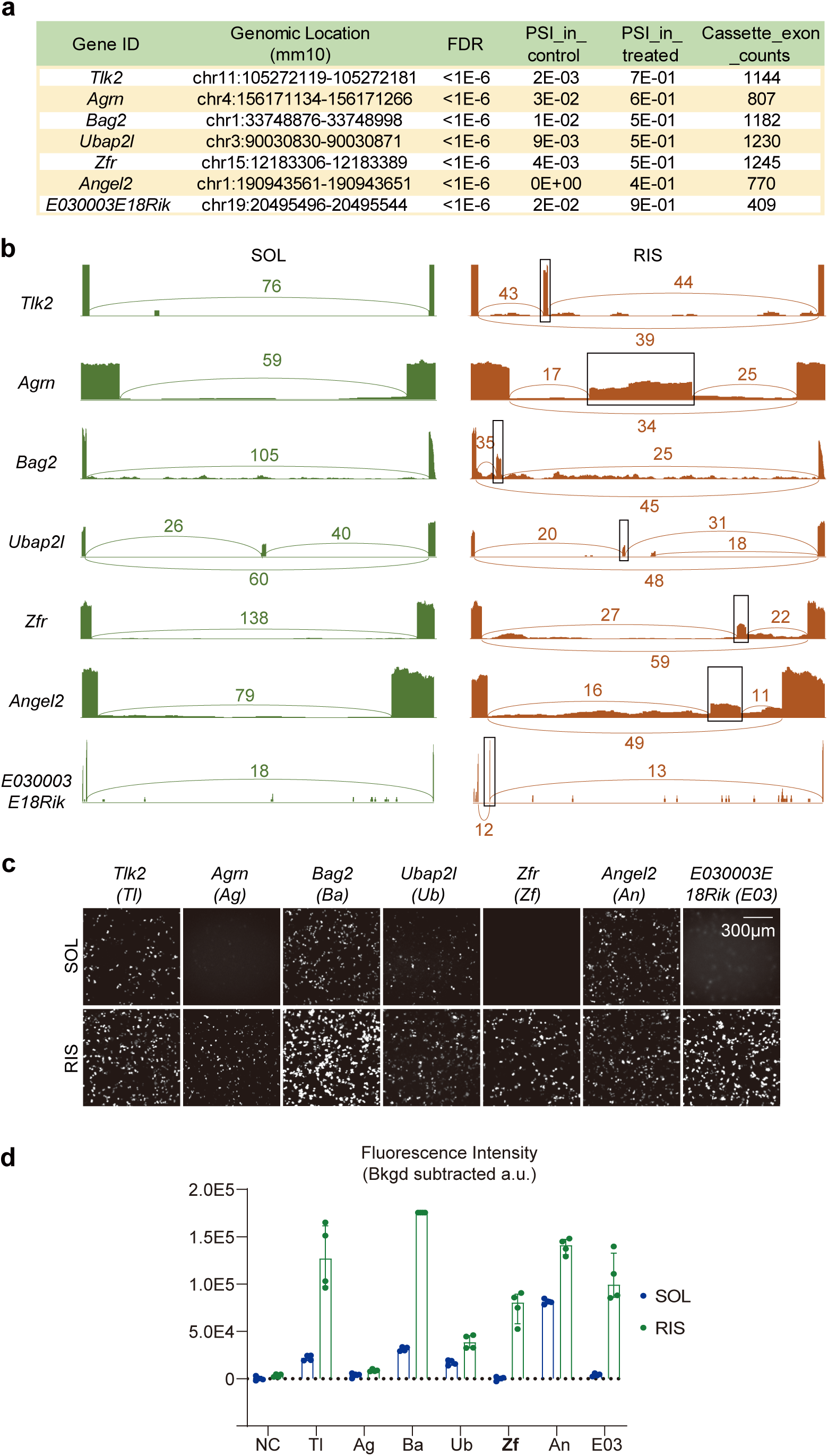
**Screen for candidates of risdiplam-induced alternative splicing modules**. **a**, Representative outputs of CASH analysis on RNA sequencing data. Genes are selected by FDR < 0.05, PSI < 0.05 in the control group, PSI > 0.40 in the treated group and total cassette exon counts > 400. FDR, false discovery rate using the Benjamini-Hochberg method. PSI, percent spliced in. **b**, Sashimi plots of the candidate modules in the genome. Pseudoexons enclosed in the box. **c-d**, Representative fluorescence images of GFP signals in HEK293T cells treated with reporters controlled by the candidate modules. Fluorescence signals were quantified by a microplate reader. NC, negative control. SOL, solvent. RIS, risdiplam. Bkgd, background. N = 4 biological repeats per group.

**Extended Data Fig. 2:**
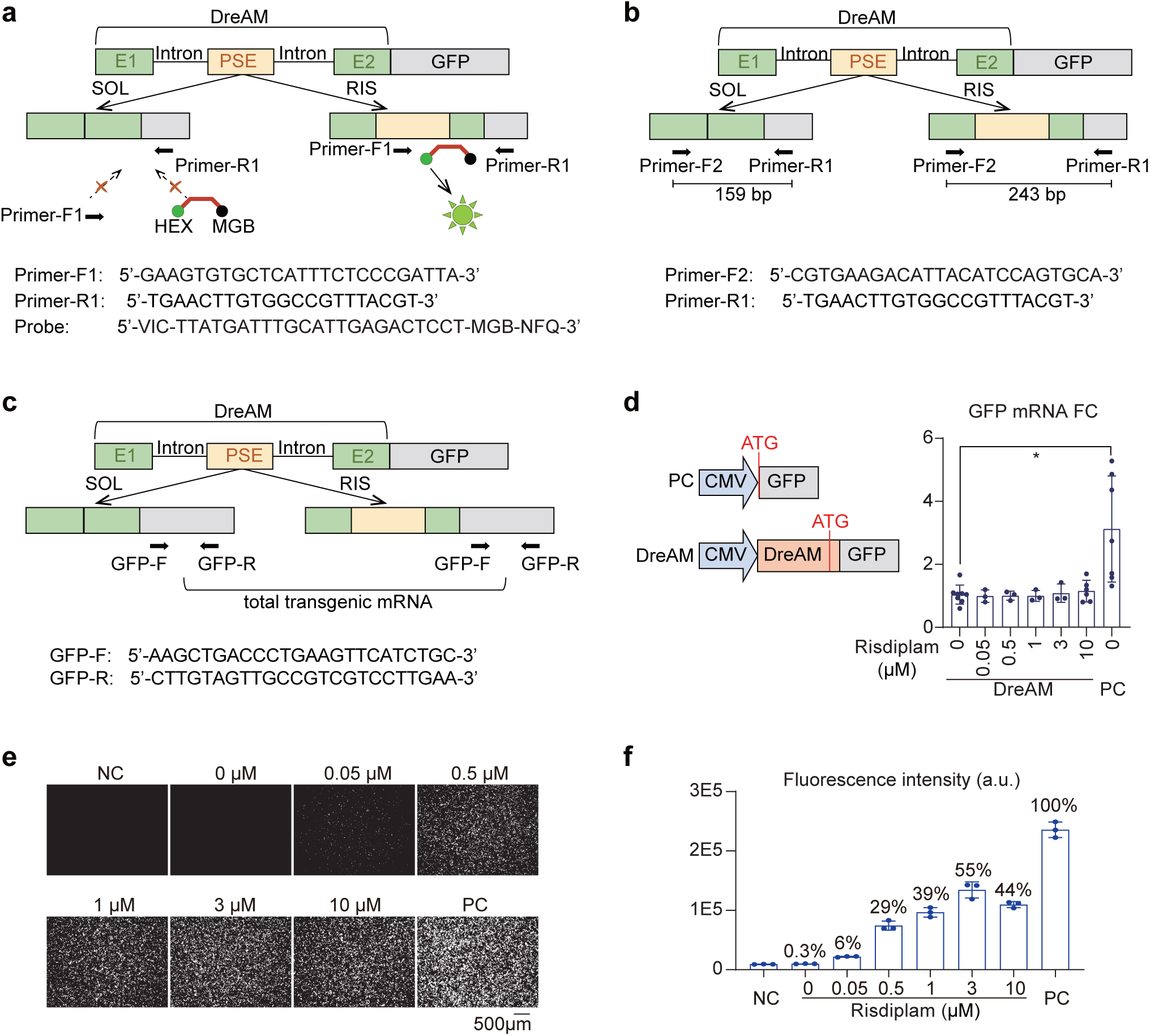
**Characterization of DreAM activity in HEK293T cells**. **a**, Design of a TaqMan-based RT-qPCR assay to specifically measure the PSE-included mRNA. The TaqMan probe targets the sequence spanning the PSE-exon 2 junction. **b,** Design of an RT-PCR assay to measure the ratio of RNA with or without PSE spliced-in. **c,** Design of an RT-qPCR assay to quantify total mRNA regardless of the PSE splicing states. **d,** RT-qPCR quantification of total GFP mRNA after plasmid transfection for 24 h. Student’s t-test, *p < 0.05. **e-f,** Fluorescence images and quantification of GFP intensity by a microplate reader. GFP signal relative to PC is shown above the column in **f**. NC, negative control. PC, positive control. N = 3 biological repeats. a.u., arbitrary unit.

**Extended Data Fig. 3:**
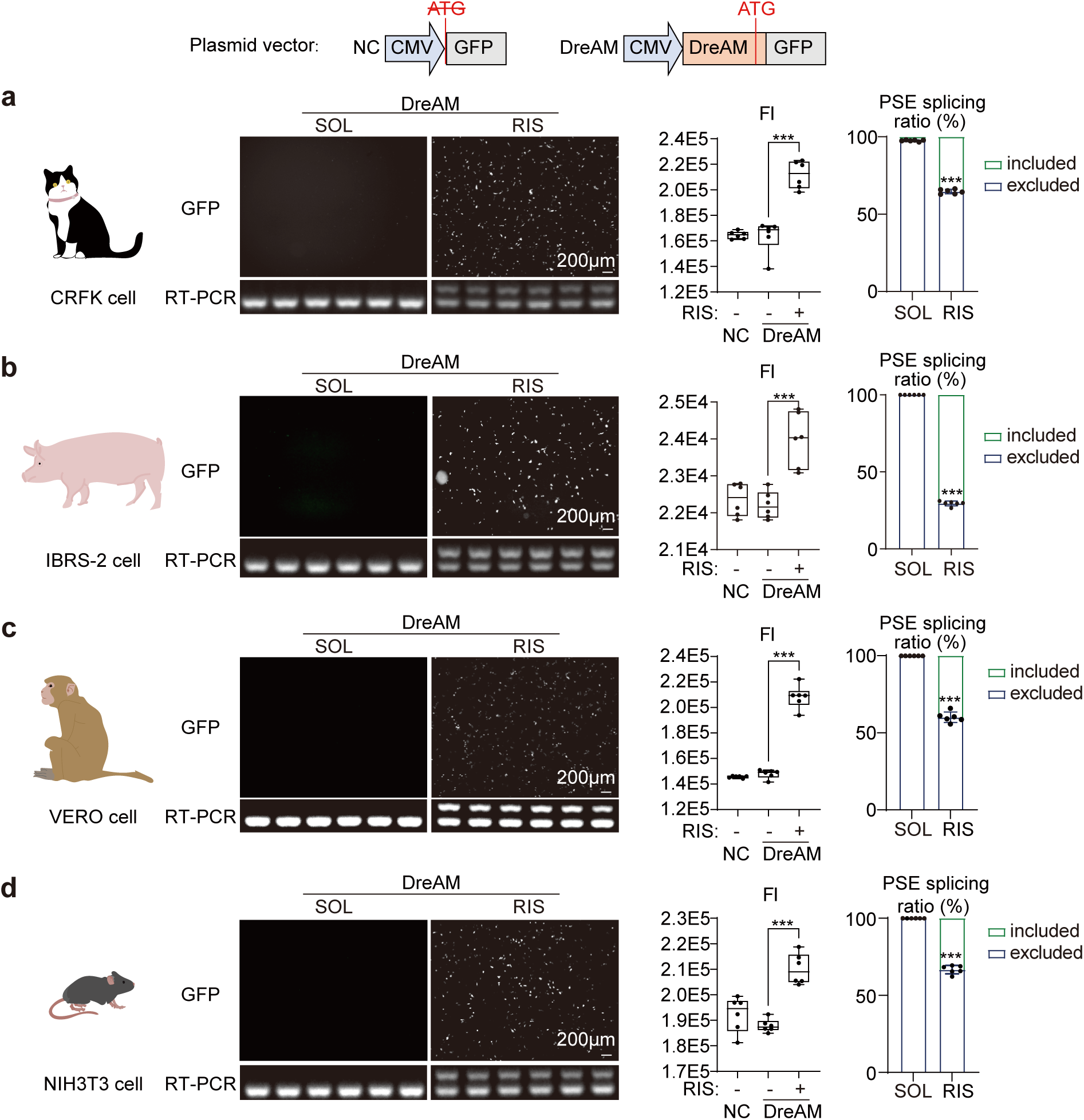
**Characterization of DreAM activity in different species**. DreAM-GFP reporter plasmids were transfected for 24h before risdiplam (RIS) or solvent (SOL) was added for another 24 h. Cells were imaged under a fluorescence microscope, fluorescence intensity quantified by a microplate reader. RNA was then extracted for RT-PCR analysis. These analyses were performed in the following cell lines: **a**, CRFK, Crandell-Rees Feline Kidney cells from cat. **b,** IBRS-2, Instituto Biologico-Rim Suino-2 cells from pig. **c,** VERO cells from African green monkey. **d**, NIH3T3, an immortalized murine fibroblast cell line. N = 6 biological repeats per group. Student’s t-test, ***p<0.001.

**Extended Data Fig. 4:**
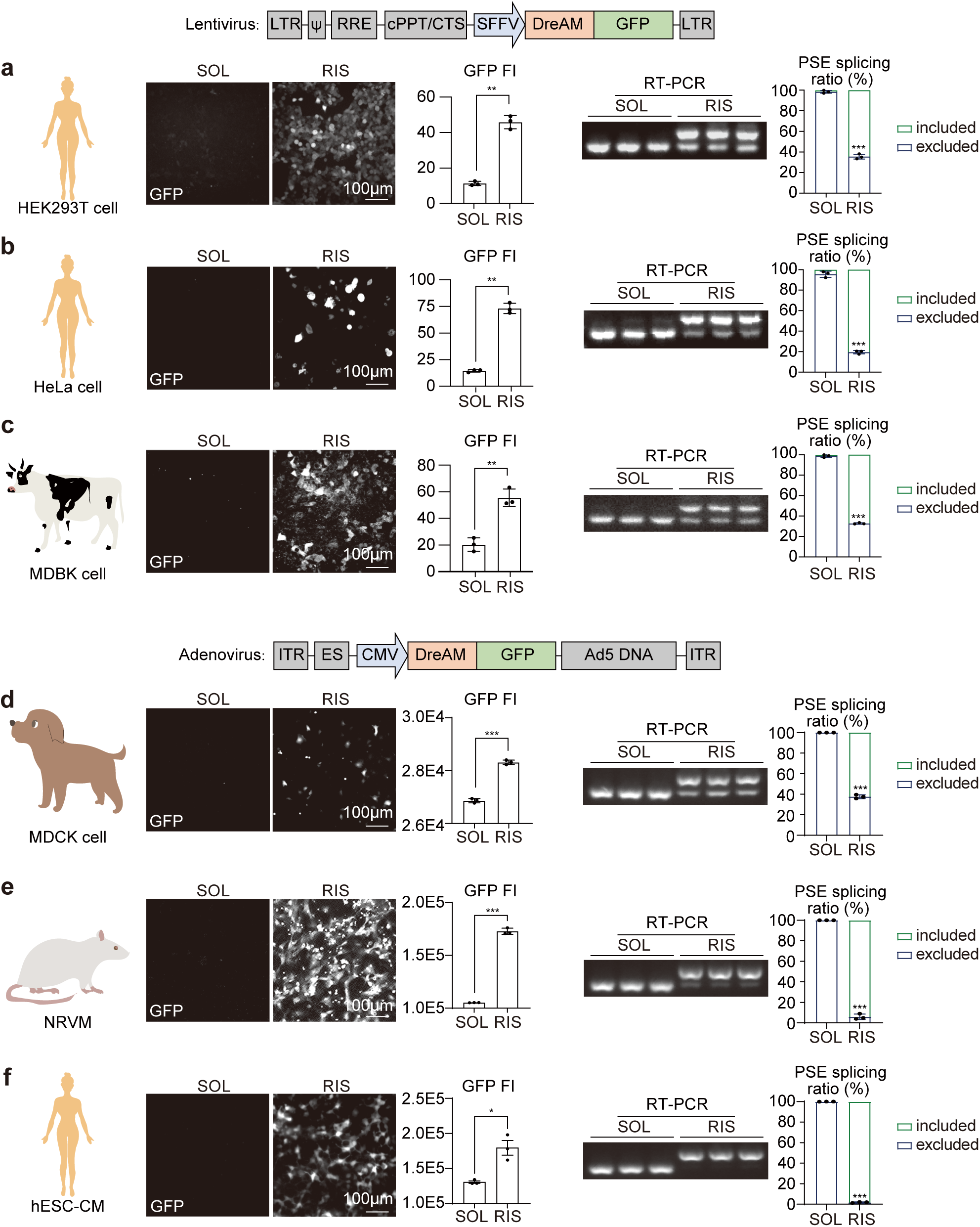
**Characterization of DreAM activity through different vectors**. The DreAM-GFP reporter was packaged into lentivirus **(a-c)** and adenovirus **(d-f)**, which were transduced into the following cell lines for fluorescence and RT-PCR analysis after risdiplam (RIS) or solvent (SOL) treatment. **a**, HEK293T cells derived from human embryonic kidney. **b,** HeLa cells from human cervical cancer. **c,** MDBK, Madin-Darby bovine kidney cells. **d**, MDCK, Madin-Darby canine kidney cells. **e**, NRVM, neonatal rat ventricular cardiomyocytes. **f**, hESC-CM, human embryonic stem cell-derived cardiomyocytes. In **a-c**, fluorescence intensity (FI) was quantified by ImageJ. In **d-f**, fluorescence intensity (FI) was measured using a microplate reader. N = 3 biological repeats. Student’s t-test, *p<0.05, **p<0.01, ***p<0.001.

**Extended Data Fig. 5:**
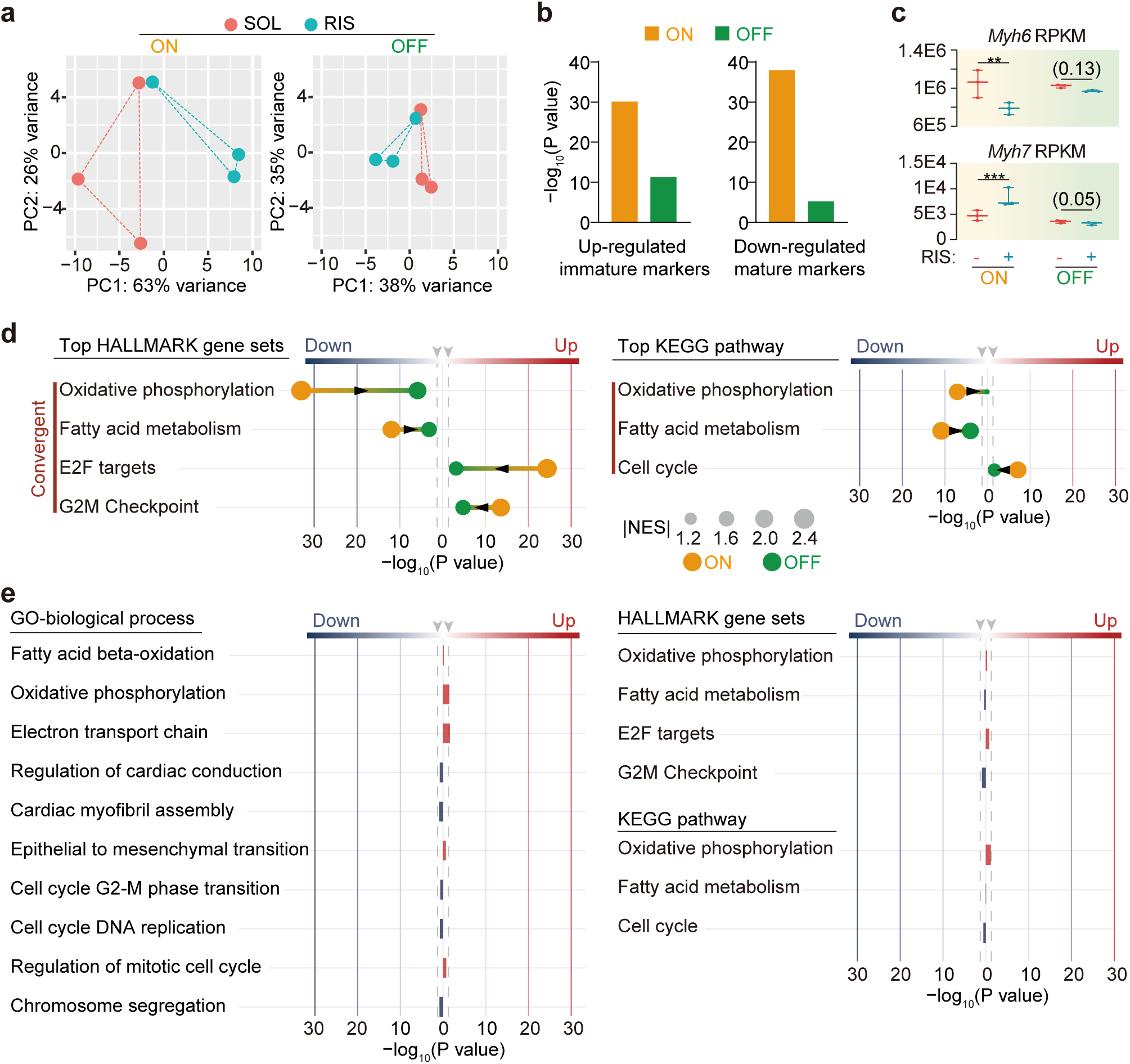
**Convergent gene expression changes at ON and OFF stages**. **a**, Principal component analysis (PCA) of RNA seq data at ON and OFF timepoints. **b,** GSEA using gene sets representing immature (P7) and mature (P30) cardiomyocytes. NES, normalized enrichment score. **c,** *Myh6* and *Myh7* RNA-Seq quantification. RPKM, Reads Per Kilobase per Million mapped reads. DESeq2-based statistical analysis, **p<0.01, ***p<0.001. Non-significant p value in parenthesis. **d,** GSEA of representative HALLMARK and KEGG gene sets at ON and OFF timepoints. KEGG, Kyoto Encyclopedia of Genes and Genomes. **e,** GSEA of the same gene sets as shown previously using RNA-seq data about risdiplam treatment alone in the absence of AAV.

**Extended Data Fig. 6:**
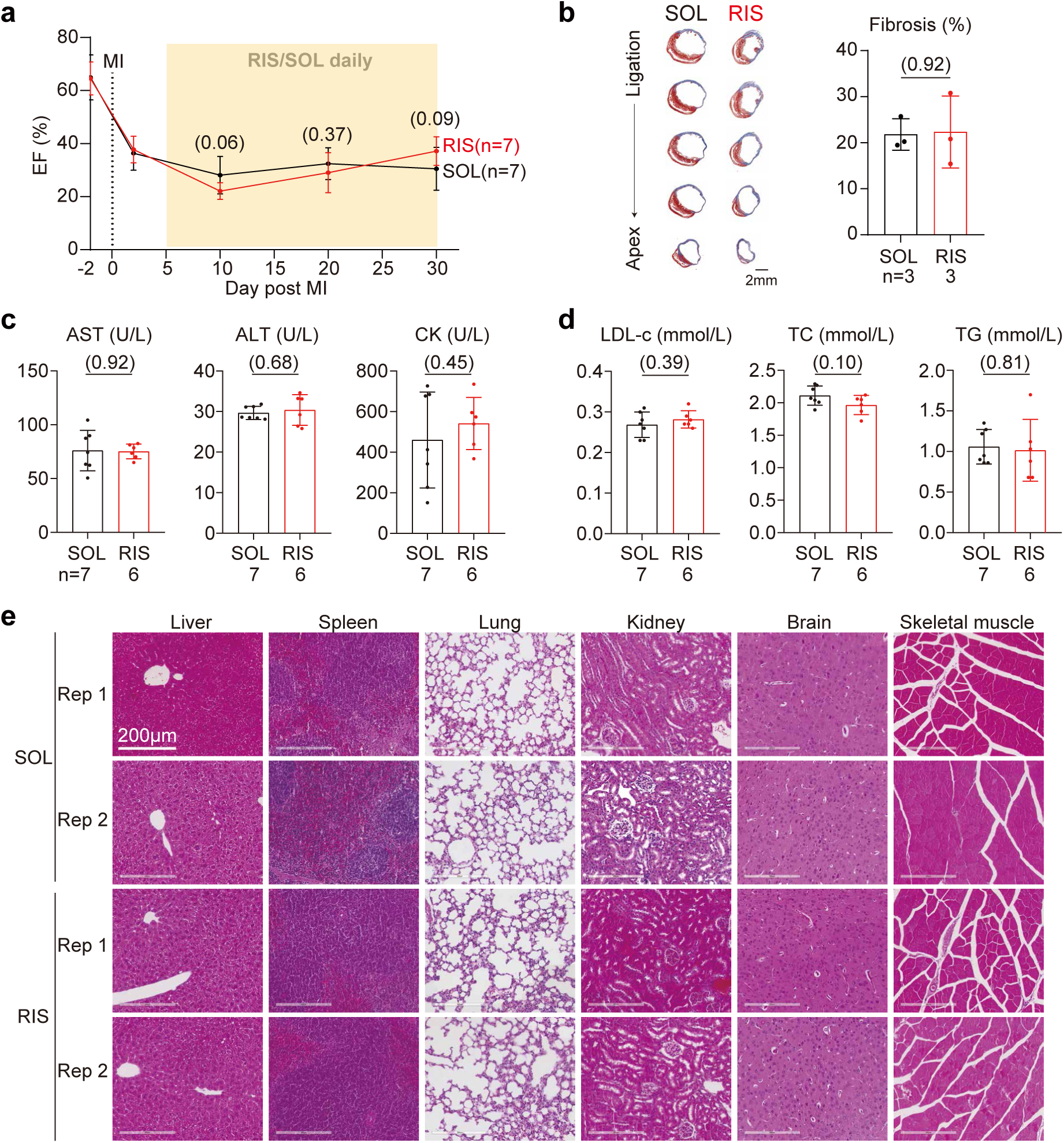
**The impact of risdiplam on myocardial infarction in mice**. **a,** Ejection fraction of MI mice undergoing daily 10 mg/kg risdiplam or solvent treatment. **b,** Masson’s trichrome staining and quantification of fibrosis on heart sections from risdiplam/solvent-treated mice at day 30 post MI. **c-d,** Serum biomarker levels at day 30 post MI. AST, aspartate aminotransferase. ALT, alanine aminotransferase. CK, creatine kinase. LDL-c, low-density lipoprotein cholesterol. TC, total cholesterol. TG, triglyceride. Student’s t test, non-significant p value in parenthesis. Mean ± SD. **e,** Hematoxylin and eosin staining of major organs at one month after MI. Rep, replicate.

**Extended Data Fig. 7:**
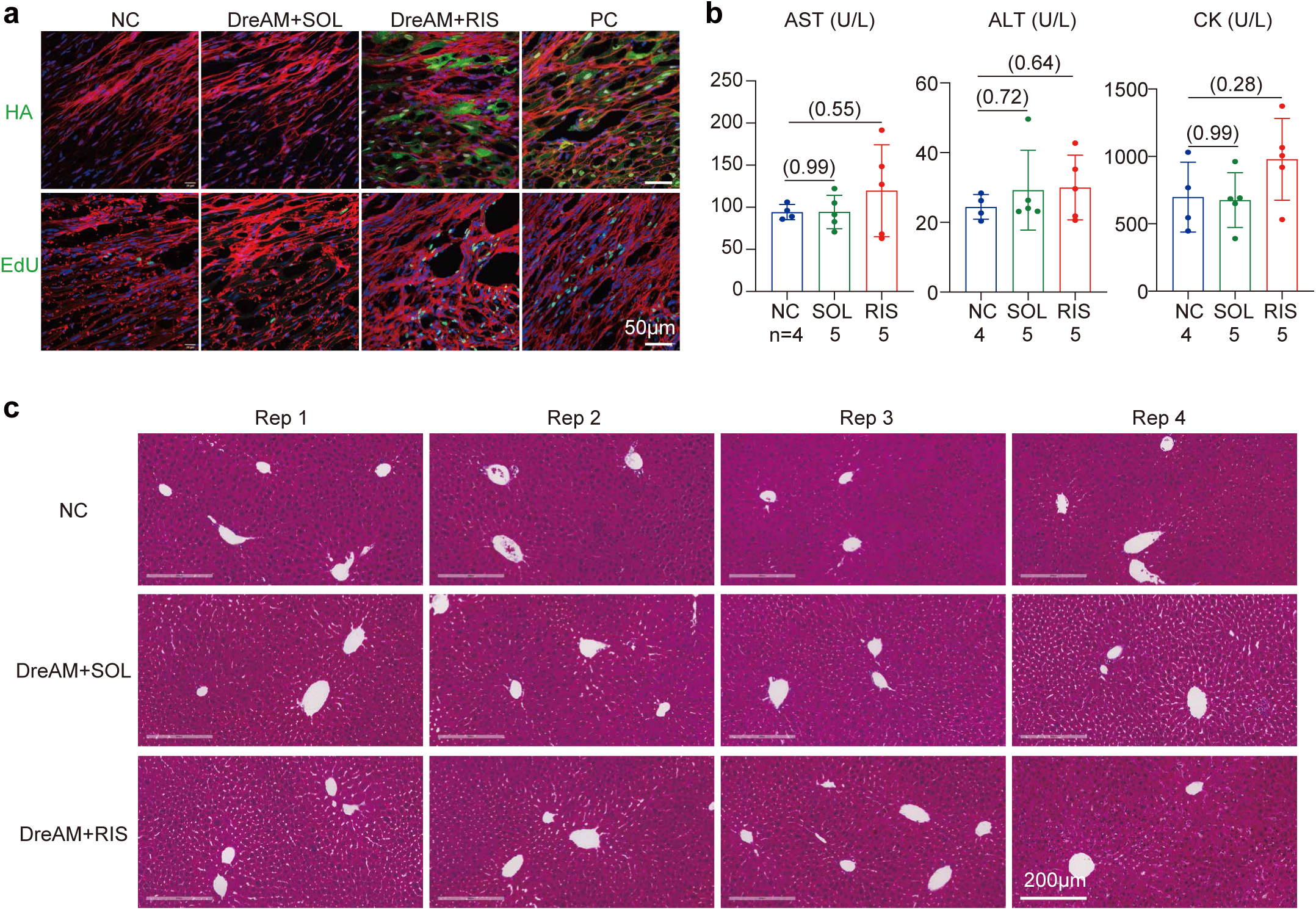
**DreAM-controlled YAP^5SA^ expression and the impact on liver**. **a,** Fluorescence images of HA and EdU in the border zone of MI hearts. NC, negative control. PC, positive control. DreAM+SOL, AAV-DreAM-YAP^5SA^ followed by solvent treatment. DreAM+RIS, AAV-DreAM-YAP^5SA^ followed by risdiplam treatment. Samples were collected one day after the last risdiplam/solvent treatment. **b,** Serum AST, ALT and CK levels at one month post MI. AST, aspartate aminotransferase. ALT, alanine aminotransferase. CK, creatine kinase. **c,** Hematoxylin and eosin staining of liver samples at one month after MI. Rep, replicate.

## Notes

### Competing Interest Statement

The authors have declared no competing interest.

