## Supplementary material for "A Drug-Elicitable Alternative-Splicing Module (DreAM) for Tunable AAV Expression and Controlled Myocardial Regeneration": Methods, Supplementary_Data_1∼7, Supplementary_Table1∼4: Supplementary-Methods.docx

Viral production

AAV production was performed by PackGene Biotech. In brief, 140 μg AAV-ITR, 140 μg AAV-Rep/Cap and 320 μg pHelper (pAd-deltaF6, Penn Vector Core) plasmids were produced by QIAGEN Plasmid Kits (12165, Qiagen) and transfected into 10 15-cm plates of HEK293T cells using PEI transfection reagent (40816ES03, Yeasen) as previously described^37^. 60 hours after transfection, cells were scraped out, resuspended in PBS and lysed by 4 freeze-thaw-vortex cycles. Cell debris were removed after 10,000g centrifugation. AAV in cell culture medium was precipitated by 8% PEG8000 (P2139, Sigma-Aldrich), resuspended in PBS, and pooled with cell lysates. AAV was purified in a density gradient (D1556-250ML, Sigma) by ultracentrifugation and concentrated in PBS using an ultrafiltration tube (100K molecular weight cutoff, Millipore). AAV titer was absolutely quantified by real-time quantitative PCR using primers amplifying a fragment of ITR.

Lentivirus production was performed by PackGene Biotech. In brief, psPAX2, pMD2.G and LV3-SFFV-DreAM-GFP were produced by QIAGEN Plasmid Kits (12165, Qiagen). Triple transfection was performed via the PEI (40816ES03, Yeasen) method into a 10-cm plates of HEK293T cells. Virus was harvested at 48h, 72h, and 96h post transfection from the cell culture medium after centrifugation at 880g for 5 minutes. The lentivirus was then purified by ultracentrifugation, concentrated using an ultrafiltration tube. Lentivirus titer was absolutely quantified by real-time quantitative PCR using primers amplifying a fragment of ITR.

Adenovirus production was performed by Hanbio. In brief, 4 μg of recombinant adenoviral plasmids were purified by Plasmid DNA purification kits (740412, MACHEREY-NAGEL) and transfected into a 6-cm plates of HEK293A cells using 15 μl LipoFiter transfection reagent (HB-TRCF-1000, Hanbio). After 3 serial cell passages, cells and supernatant were harvested and virus were released by 4 sequential freeze–thaw cycles. The supernatant was collected after the lysates were centrifuged at 500g for 8 minutes. Then the adenovirus was enriched by centrifugation at 100,000g and dialyzed into PBS for 3 times before storage. The adenovirus titer was measured via the TCID50 assay.

Tissue sectioning and histology analysis

Hearts were fixed by perfusion on a Langendoff system with 4% PFA for 15 minutes. Alternatively, hearts and other tissues were immersed in 4% PFA at 25°C for 2-4 hours. Fixed tissues were dehydrated in sucrose solutions (15% followed by 30%) overnight. For frozen sectioning, the tissue samples were embedded in OCT (Sakura) and frozen in -20°C overnight. Sections (7 μm) were cut on a cryostat microtome (CM1950, Leica).

For paraffin sectioning, the sucrose dehydrated samples were further dehydrated through a gradient of ethanol and N-butanol (70% and 80% ethanol for 3 hours; 1:1 90% and N-butanol solution overnight; 1:1 95% and N-butanol twice for 45 minutes each; N-butanol for 45 minutes once and 15 minutes once). Then the samples were waxed in liquid paraffin for 2 hours and embedded to make paraffin blocks using a tissue embedder (EG1150, Leica, Germany). The tissues were cut into 4 μm sections by a paraffin slicer (RM2245, Leica, Germany) and adhered to the slides on a water bath (HI1210, Leica, Germany). Being dried at 65°C for 30 minutes, the slides were deparaffined through the following solutions: in xylene for 3 times, 10 minutes each; in anhydrous ethanol for twice, 5 minutes each; and in 95%, 90%, 80%, 70% ethanol once for 5 minutes each. After being washed with PBS for 3 × 5 minutes, the sections were used for staining.

Hematoxylin-Eosin staining, Masson's trichrome staining and Picrosirius red staining were performed by Wuhan Servicebio Technology. Hematoxylin-Eosin staining was performed using the Hematoxylin-Eosin (H&E) HD kit (G1076, Servicebio) according to the instructions. The sections were stained in the HD constant staining pretreatment solution for 1 min then in Hematoxylin solution for 3-5 min. After being rinsed with tap water, the sections were stained in Hematoxylin Differentiation solution for 3-5 s, rinsed with tap water, stained in Hematoxylin Bluing solution for 3-5 s and then rinsed with tap water. After rinsing in 95% ethanol for 1 min and Eosin dye for 15 s, the sections were dehydrated in absolute ethanol for 3 times, normal butanol for 2 times, xylene for 2 times before sealing with neutral balsam.

For Masson's trichrome staining, the sections were stained with Masson dye solution set (G1006, Servicebio) following the manual. Briefly, the slices were soaked in Masson A overnight then rinsed with tap water. Masson B and Masson C were mixed into Masson solution at 1:1 ratio. Then the sections were put into Masson solution for 1 min, differentiated with 1% hydrochloric acid alcohol for several seconds, rinsed with tap water, soaked in Masson D for 6 min, rinsed with tap water, soaked in Masson E for 1 min and Masson F for 2-30s. The sections were rinsed with 1% glacial acetic acid and then dehydrated with anhydrous ethanol for 2 times. For clearing and sealing, slides were soaked in 100% ethanol for 5 min followed by xylene for 5 min before sealing with neutral balsam.

For Picrosirius Red staining, the slices were stained in Picrosirius Red staining solution (G1018, Servicebio) for 8 min and then dehydrated quickly with anhydrous ethanol for three times. Lastly, the sections were put into xylene for 5 min before sealing with neutral balsam.

Stained sections were imaged using an OCUS 40 microscope slide scanners (Grundium). To determine the fibrotic area, the intermediate heart sections of the serial sections were measured. ImageJ was used to distinguish and measure the scar area and the whole tissue area. The scar area divided by the whole tissue area gives rise to the fibrosis area percentage.

Bioluminescence imaging

Sterile D-PBS (60152ES76, Yeasen) and D-Luciferin (40902ES03, Yeasen) were used to prepare 15 mg/ml luciferin solution, which were injected intraperitoneally 10 minutes before imaging at the luciferin/body weight ratio of 150 mg/kg. Animals were anesthetized via 3% isoflurane inhalation before bioluminescent imaging and analysis in a IVIS Spectrum In Vivo Imaging System (PerkinElmer).

Echocardiography

Echocardiography was performed on a VINNO 6 VET ultrasound system (PHYSIA). Echocardiography was performed by technicians blinded to the groups. Animals were anesthetized via 3% isoflurane before ultrasound imaging. The left ventricle internal diameters, left ventricle posterior wall thickness, and ejection fraction were evaluated by M-mode echocardiography.

Blood test

Blood was allowed to clot for 30 min at room temperature. After centrifugation at 6000 rpm for 10 minutes, serum was obtained from the supernatant. Serum components were measured using a chemistry analyzer (BS-430, Mindray) using kits following manufacturer’s instructions (Mindray, Shenzhen, China). Aspartate aminotransferase (AST, V276176), alanine aminotransferase (ALT, V501534) and creatine kinase (CK, V251576) were measured following the IFCC methods. LDL-Cholesterol (LDL-c) was measured using a direct method (V251566). Total Cholesterol (TC, V501668) and triglycerides (TG, V501666) was measured via the POD Method.
